# Cross-species spatial profiling links progranulin to an immune suppressive niche in brain metastasis

**DOI:** 10.64898/2026.09.13.751277

**Authors:** Nicole M. Eskow, Kylie Prutisto-Chang, Maria A. Gomez-Munoz, Minkyung Kang, Irineu Illa Bochaca, Milad Ibrahim, Sorin A.A. Shadaloey, Amanda Flores Yanke, Luiz Henrique Geraldo, Giuseppe Barisano, Anuja Sathe, Pablo Nunez, David A. Solomon, Elif Tugce Karasu, Hanlee Ji, Amanda Lund, Maxine Umeh Garcia, Melanie Hayden Gephart, Jun Wang, Eva Hernando

## Abstract

Immune checkpoint inhibitors (ICI) have revolutionized the treatment of brain metastasis (BM), demonstrating intracranial response rates and durability not previously seen with other therapies. However, approximately 50% of patients do not respond to ICI, and the brain-specific interactions that shape anti-tumor immunity remain poorly understood. We profiled five syngeneic BM models using spatial transcriptomics, identifying myeloid-rich niches enriched for interferon-responsive and disease-associated microglia (DAM)-like programs. Ligand-receptor colocalization analyses identified progranulin (PGRN) as a candidate mediator of these niches. Host- or tumor-cell *Grn* loss reduced BM burden, and *Grn*-deficient macrophages exhibited altered metabolic programs and tumor-cell engulfment in vitro. Across independent human BM datasets, *GRN* expression was associated with conserved lysosomal and DAM-like myeloid programs, and *GRN*-high myeloid regions colocalized with immunosuppressive signatures and dysfunctional CD8+ T cell states. These cross-species findings identify PGRN as a candidate mediator of the BM immune niche and support further investigation of its therapeutic relevance.

**Highlights:**

- A multi-model spatial atlas resolves myeloid niches in both preclinical models of brain metastases and human patient tissues
- Host- and tumor-cell progranulin support brain metastatic growth
- Progranulin associates with lysosomal function and disease-associated myeloid states in brain metastasis
- Human progranulin-rich myeloid niches colocalize with dysfunctional cytotoxic T cells in brain metastasis

**Graphical Abstract:** 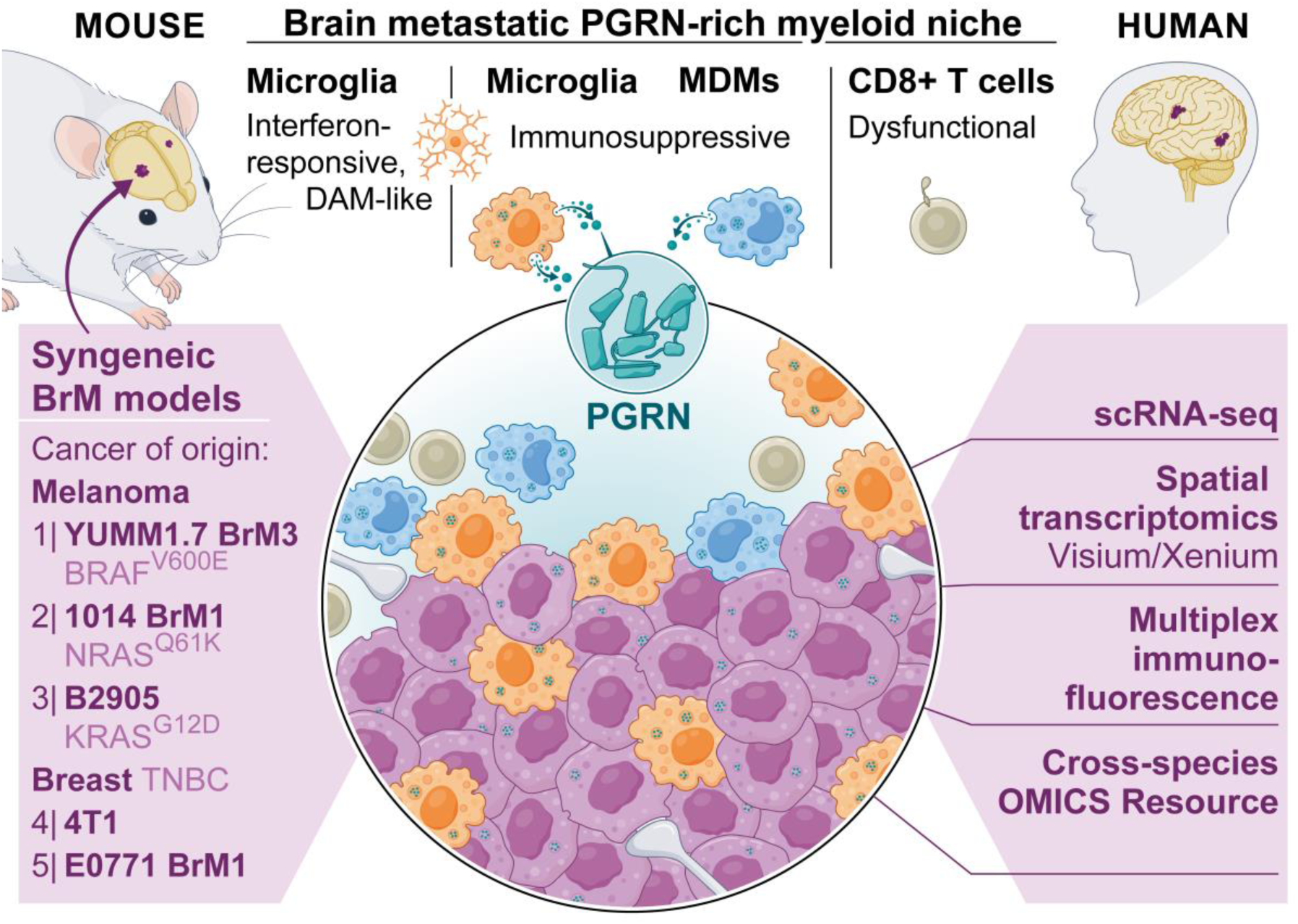

## Introduction

Brain metastases (BM) are the most common malignant intracranial tumors in adults and are a major cause of morbidity and mortality in patients with advanced cancer. Lung cancer, breast cancer, and melanoma account for up to 80% of cases, and the incidence of BM has increased with improved detection and increases in survival of patients with extracranial disease ^1–3^. Although systemic therapies historically have demonstrated limited intracranial activity, advances in targeted therapy and immunotherapy have expanded treatment options for patients with BM^4^. In melanoma BM, combined PD-1/CTLA-4 blockade has produced durable intracranial responses, particularly in patients with asymptomatic disease^5,6^. Nevertheless, many patients are intrinsically resistant to immune checkpoint inhibition (ICI), and clinical benefit remains limited among patients with symptomatic disease or corticosteroid dependence. Similarly, many initial ICI responders develop recurrent, resistant disease^1^. These limitations underscore the need to identify additional immune mechanisms that could be therapeutically targeted either alone or in combination with existing treatment regimens.

The BM microenvironment is composed of tumor cells, tissue-resident central nervous system populations, and recruited immune and stromal cells. Single-cell studies of human BM have consistently identified a prominent and heterogeneous myeloid compartment containing both tissue-resident microglia and recruited monocyte-derived macrophages (MDM)^7–12^. Compared with extracranial metastases (ECM), BMs exhibit greater macrophage infiltration, with BM-associated macrophages expressing distinct immunomodulatory programs enriched for genes associated with tumor-supportive functions^7^. These BM-associated myeloid populations adopt diverse transcriptional states and may either support or restrain tumor progression depending on their cellular and molecular context. Their abundance and functional plasticity have therefore motivated efforts to therapeutically target tumor-associated microglia and macrophages (TAMs). In preclinical BM models, inhibition of CSF1/CSF1R signaling can suppress tumor growth, but adaptive myeloid responses, including compensatory CSF2 signaling, limit the durability of this effect^13^. Beyond CSF1R inhibition, targeting the CD47-SIRPα antiphagocytic axis or TREM2-associated myeloid programs has enhanced anti-tumor responses in preclinical cancer models^14,15^. In BM specifically, TREM2-expressing macrophages have been implicated in local T cell suppression^16^. Despite these promising preclinical findings, clinically effective myeloid-directed therapies for BM have not yet been established. Moreover, although single-cell studies have defined the heterogeneity of BM-associated myeloid populations, the spatial organization and intercellular signaling networks that shape these states remain incompletely understood.

In this study, we established a panel of syngeneic mouse models of BM and used spatial transcriptomics to interrogate tumor-immune and tumor-stroma interactions within the BM microenvironment. Broadly, BMs elicited robust microglial and astrocytic responses that shared transcriptional features with programs described in neurodegenerative diseases. Further analysis revealed that two microglial subtypes, an interferon-enriched inflammatory population, and a disease-associated microglia (DAM)-like population, are enriched within the BM microenvironment. Gene set enrichment and ligand-receptor interaction analyses facilitated the discovery of progranulin (PGRN, *Grn*) as a potential regulator of BM development. Progranulin is a secreted 88-kDa glycoprotein with well-described roles in microglial inflammation and lysosomal biology, and PGRN loss of function has been linked to frontotemporal dementia (FTD) and neuronal ceroid lipofuscinosis. In our models, BM burden was reduced upon silencing host and tumor-cell *Grn*, implicating both stromal- and tumor-derived PGRN in BM progression. Across mouse and human datasets, *GRN* expression was consistently associated with DAM-like myeloid programs, including lysosomal, phagocytic, and antigen-processing functions.

Orthogonal spatial transcriptomic and multiplex immunofluorescence analyses in human BM of various origins (e.g. melanoma, breast cancer) demonstrate that *GRN* is robustly expressed and co-localizes with immunosuppressive signatures and dysfunctional CD8+ T-cell states. Together, these findings identify PGRN as a candidate mediator of the BM immunosuppressive niche and support further investigation of its therapeutic potential.

## Results

### A spatial transcriptomic atlas of preclinical brain metastasis models reveals an immunologically active, myeloid-enriched microenvironment

To gain insight into the unique BM tumor immune microenvironment, we developed a panel of five murine-derived melanoma and breast cancer cell lines that exhibit high tropism (≥90% penetrance) to the brain upon intracardiac injection^17^ (Table 1, Figure S1A-B). Three of these cell lines, Yumm1.7 GFP-Luciferase (GL) BrM3, E0771 GL BrM1, and 1014 BrM1, required serial injection via either intracardiac or intracarotid instillation to derive brain-metastatic lines with high penetrance upon intracardiac injection^18,19^ (Figure S1A-B). 4T1 and B2905 cells demonstrated high brain tropism upon intracardiac injection without the need for additional in vivo selection. To comprehensively and anatomically characterize the tumor immune landscape across our murine syngeneic brain-metastatic preclinical models, we chose the 10x Genomics Visium Spatial Transcriptomics platform to perform unbiased spatial gene expression profiling. Tumor-bearing brains (1-2 per cell line) were profiled alongside sham-control brains, which provided a non-tumor reference for identifying BM-associated transcriptional programs and changes in inferred intercellular signaling (Figure 1A).

**Figure 1:**
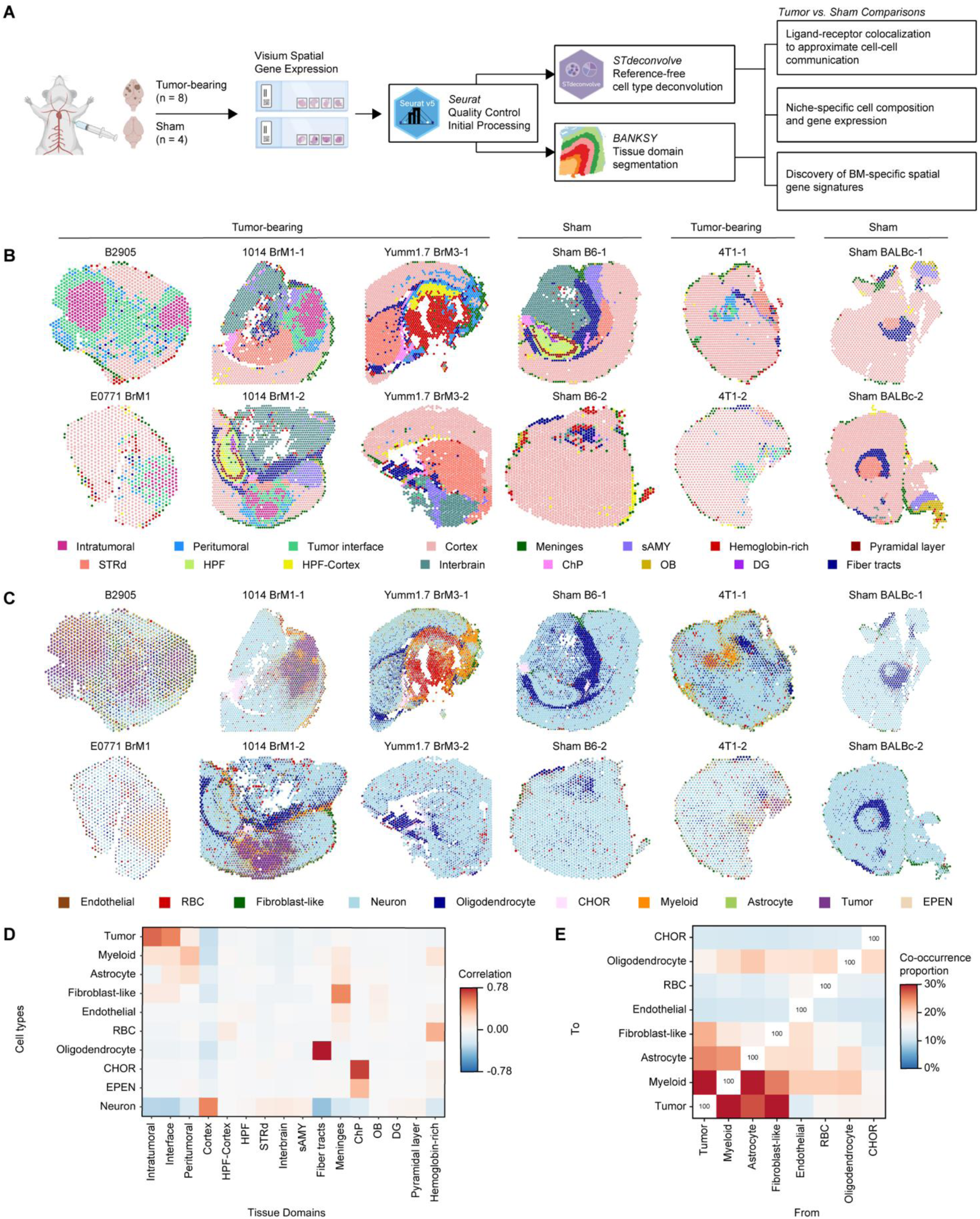
Spatial transcriptomic profiling defines the cellular and anatomic organization of the brain metastatic niche. A) Overview of the spatial transcriptomic workflow. Eight tumor-bearing brains representing five murine syngeneic brain metastasis models and four sham (PBS-injected) brains were profiled using the 10x Genomics Visium Spatial Gene Expression platform. Seurat, BANKSY, and STdeconvolve were used for initial processing, tissue-domain segmentation, and reference-free cell-type deconvolution, respectively. Figure generated in BioRender. B) Spatial maps of tissue-domain assignments across tumor-bearing and sham samples. HPF, hippocampal formation; STRd, dorsal striatum; sAMY, striatum-like amygdalar nuclei; ChP, choroid plexus; OB, olfactory bulb; DG, dentate gyrus. C) Estimated cell-type proportions per spatial transcriptomic spot obtained by LDA-based reference-free deconvolution. RBC, red blood cell; CHOR, choroid plexus-derived cells; EPEN, ependymal cells. D) Correlation between deconvolved cell-type proportions and tissue-domain assignments. E) Pairwise spot-level co-occurrence of deconvolved non-neuronal cell types in tumor-bearing brains, shown as the percentage of spots in which each pair of cell types was jointly detected.

**Table 1:** Syngeneic models of brain metastasis.

| Cell lines | Cancer of origin | Mutational status | Time to humane endpoint | Incidence of Metastasis Upon Intracardiac Injection |  |  |
| --- | --- | --- | --- | --- | --- | --- |
|  |  |  |  | *Brain* | *Liver* | *Kidney/Adrenal* |
| Yumm1.7 GL BrM3 | Melanoma | *Braf*<sup>V600E</sup> | 2 weeks | 100% | 0% | 0% |
| 1014 BrM1 | Melanoma | *Nras*<sup>Q61K</sup> | 6 weeks | 95% | 100% | 100% |
| B2905 | Melanoma | *Kras*<sup>G12D</sup> | 3 weeks | 100% | 30% | 70% |
| 4T1 GL | Breast | TNBC | 2 weeks | 100% | 50% | 100% |
| E0771 GL BrM1 | Breast | TNBC | 3 weeks | 90% | 30% | 100% |

We first applied BANKSY to define spatial domains using spot-level gene expression and neighboring transcriptomic context, followed by Harmony integration to align transcriptionally related spots across samples^20,21^(Figures 1B, S2A). Tumor-bearing regions were subsequently annotated using Seurat’s FindAllMarkers function in conjunction with manual inspection of H&E-stained tissue sections^22^ (Figure S1C). Intratumoral domains were defined as clusters located within the tumor core that expressed tumor-specific markers (e.g., *Melana* in melanocytic melanoma models, *Mgp* in Yumm1.7 BrM3, and *Krt8* in breast cancer models). Interface domains were defined as clusters adjacent to the tumor border that retained expression of tumor-specific markers, whereas peritumoral domains lacked tumor-specific markers but were still transcriptionally distinct from the surrounding tumor-free brain regions. (Figures 1B, S2A-B).

To infer cell type composition from Visium spot-level transcriptomic data, we applied a reference-free deconvolution approach using Latent Dirichlet Allocation (LDA), implemented via the R package STdeconvolve^23^. Leveraging markers and expression profiles from previously annotated single cell-level datasets of mouse brains, we annotated ten broad cellular populations in our samples, including tumor cells, myeloid-lineage cells, astrocytes, neurons, oligodendrocytes, endothelial cells, fibroblast-like cells, choroid-plexus derived cells (CHOR), ependymal cells (EPEN) and red blood cells (RBCs)^24^ (Figure 1C). To independently evaluate our spatial domain annotations, we compared inferred cell-type proportions across annotated regions. Tumoral domains were enriched for tumor cells, myeloid cells, astrocytes, and fibroblast-like cells, providing additional support for the biological relevance of the domain assignments (Figure 1D). Consistent with these findings, colocalization analysis demonstrated that tumor cells most frequently co-occurred with myeloid cells within individual Visium spots (Figure 1E). Finally, independent examination of established myeloid (*Aif1, Cd68, Trem2,* and *Lyz2*) and astrocyte (*Gfap, Aqp4,* and *Serpina3n*) markers, together with model-specific tumor-marker expression, demonstrated that myeloid-associated expression was concentrated within tumor-bearing regions^24–26^ (Figure S2C). Together, these complementary analyses validate myeloid cells as a prominent component of the BM microenvironment.

Although LDA-based spot-level deconvolution can provide useful estimates of cellular composition within spatial transcriptomics data, it infers cell populations based on patterns of gene expression rather than directly identifying biologically defined cell types. Additionally, reference-free deconvolution alone cannot distinguish whether intratumoral macrophages originate from tissue-resident microglia, MDM, or a mixture of both populations. As a complementary method to deconvolve the cellular composition of the BM microenvironment, we utilized a published single-cell RNA sequencing (scRNA-seq) dataset of mouse melanoma BM to identify marker genes exclusive of immune and stromal populations^27^. By cross-referencing this dataset with previously published markers of brain and immune cell populations, we curated cell-type-specific genes signatures and mapped these onto our spatial transcriptomic samples^24,28^ (Figure 2A; Tables S1-S2). The microglia signature was enriched across the tumoral domains of all tumor-bearing samples, along with a mild enrichment of NK-cell markers. Endothelial cells (ECs) and other macrophage populations, namely MDMs, and border-associated macrophages (BAM), were enriched in the peritumoral region (Figure 2B). Unlike our independently curated astrocyte marker set (*Gfap, Aqp4,* and *Serpina3n*), which was enriched at the tumor periphery (Figure S2C), the astrocyte signature derived from the Rodriguez-Baena et al. scRNA-seq dataset did not show a clear spatial enrichment pattern in our atlas, likely because the dataset-derived signature contains broad astrocyte markers rather than those specific to reactive states. Furthermore, there was overall low detection of T cell signatures across both tumor-bearing and non-tumor-bearing regions. Together, reference-free LDA-based deconvolution and scRNA-seq-derived gene signature mapping reveal a rich myeloid infiltrate in BM, including a substantial contribution from tissue-resident microglia.

**Figure 2:**
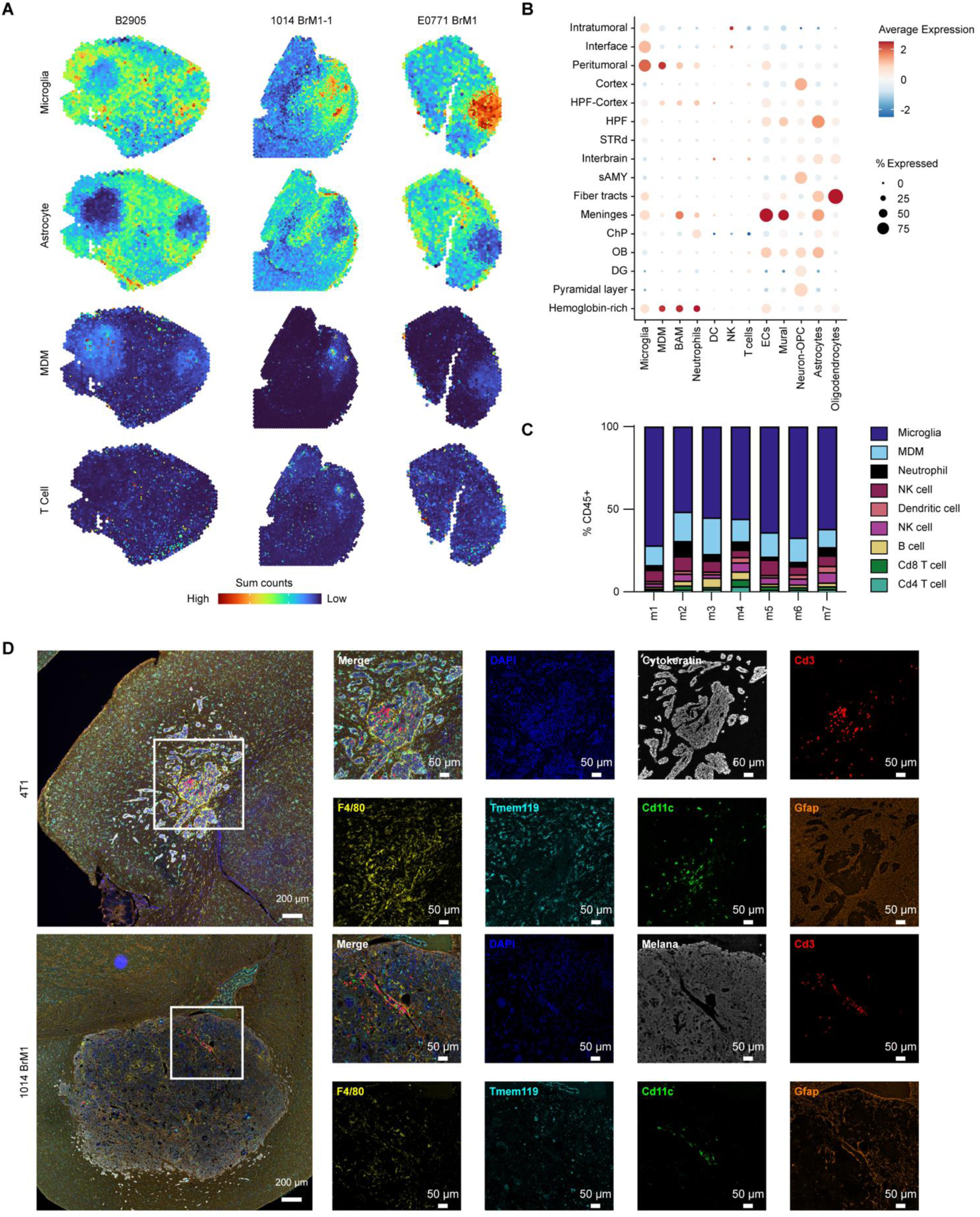
Spatial transcriptomic and protein-level profiling reveals a myeloid-rich brain metastatic niche. A) Representative spatial feature plots of microglial, astrocyte, MDM, and T cell transcriptional signatures. B) Expression of cell-type transcriptional signatures across spatial tissue domains. Neuron-OPC, mixed neuron/oligodendrocyte precursor cell signature; ECs, endothelial cells; NK, natural killer cells; DC, dendritic cells; BAM, border-associated macrophages; MDM, monocyte-derived macrophages. C) Distribution of CD45+ immune cell populations across seven B2905 tumor-bearing brains following whole brain dissociation, cell-surface marker staining, and flow cytometric analysis. D) Representative multiplex immunofluorescence staining of brain metastases. Tumor cells were identified using tumor-lineage markers, with TMEM119, F4/80, and CD11c used to characterize myeloid populations and CD3 used to identify T cells.

To validate our transcriptomic findings, we used flow cytometry to explore the immunologic compartment (CD45+ cells) of B2905 tumor-bearing cerebrums, using Cd49d as a marker to differentiate microglia (Cd49d-) from MDM (Cd49d+)^29^. In agreement with our transcriptomic analysis, microglia comprise the largest population of CD45+ cells (mean = 61.10% stdv = 7.31%), followed by MDM (mean = 15.12% stdv = 3.79%) (Figures 2C, S3A). As reflected in the spatial transcriptomics data, lymphocyte populations were present at low levels. We also performed multiplex immunofluorescence to spatially characterize the microenvironment of our BM models. We again find that microglia (TMEM119+) and MDMs (F4/80+TMEM119-) surround tumor cells, although immunofluorescence demonstrated a more prominent non-microglia macrophage population than was suggested by the transcriptomic and flow cytometry analyses. We also observed low levels of intratumoral CD3+ T cells and CD11c+F4/80-TMEM119-dendritic cells (Figures 2D, S3B-C). Taken together, the flow cytometry and immunofluorescence data corroborate the rich myeloid-lineage infiltration observed in our spatial transcriptomics data.

### Interferon-enriched and DAM-like microglial programs localize to the immediate BM microenvironment

Next, we sought to characterize the biological heterogeneity of microglia within the BM microenvironment. We subset the microglial population in the Rodriguez-Baena et al. scRNA-seq dataset and applied non-negative matrix factorization (NMF) to identify transcriptional programs underlying microglial heterogeneity^27^. NMF revealed five programs that broadly recapitulated the clusters described in the original dataset (Figure S4A, Table S3): a Homeostatic program marked by *Hpgd, Adrb2,* and *Stab1*; an Early Activation program enriched for inflammatory-response genes including *Fos, Tnf, Nfkbiz,* and *Nfkbid*; an Interferon-Enriched program characterized by canonical interferon-stimulated genes from the *Ifit* and *Oas* families; a DAM-like program enriched for phagolysosomal and lipid metabolism genes alongside established DAM markers including *Cst7, Apoe, Cd63, Itgax, Lyz2,* and *Timp2*; and a Proliferative program defined by cell cycle-related genes including *Mki67, Top2a,* and *Knl1*^30–38^. For downstream visualization and comparison, each cell was assigned to its highest-scoring NMF program to produce five microglial populations with distinct transcriptional marker genes (Figures 3A, S4B). KEGG pathway enrichment supported the program annotations, with Early Activation microglia enriched for MAPK, TNF, and cellular stress-response pathways, and DAM-like microglia enriched for phagosome formation and antigen presentation^39,40^ (Figure S4C). Pseudotime analysis, with the Homeostatic group specified as the computational root, revealed a trajectory in which Early Activation microglia occupied an intermediate position bridging the Homeostatic group to branches containing the Interferon-Enriched or DAM-like microglia (Figure 3B). This computational ordering is consistent with a model in which microglia may transition through an Early Activation state before adopting more specialized transcriptional phenotypes in BM.

**Figure 3:**
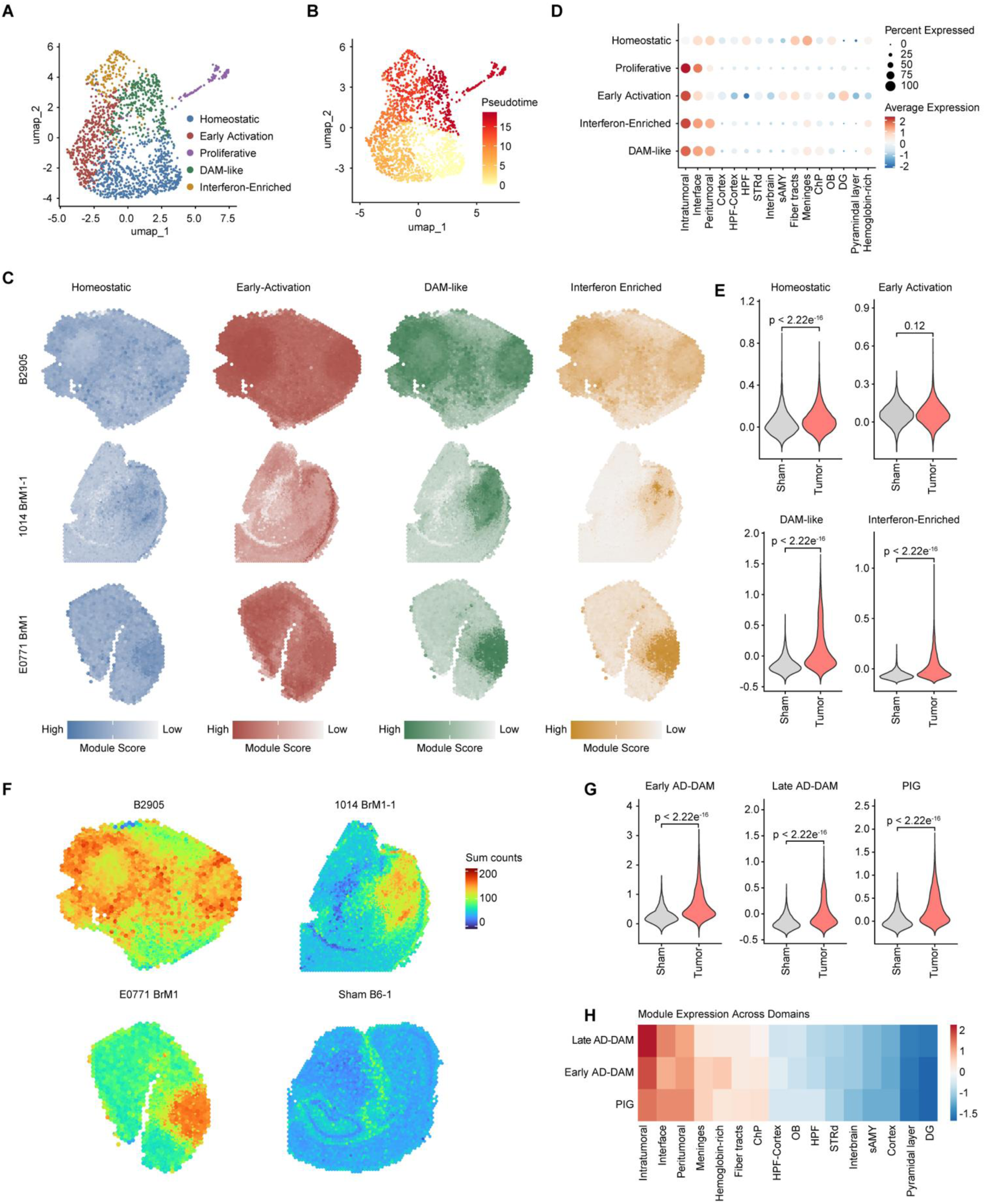
Microglia exhibit interferon-enriched and DAM-like states within the brain metastatic niche. A) UMAP projection of microglia from the Rodriguez-Baena et al. scRNA-seq dataset^27^. Cells were assigned to the NMF transcriptional program with the highest score, defining Homeostatic, Early Activation, Interferon-Enriched, DAM-like, and Proliferative states. B) Pseudotime analysis of microglia from the Rodriguez-Baena et al. dataset^27^, with the homeostatic population specified as the computational root. C) Representative spatial feature plots showing projection of Homeostatic, Early Activation, DAM-like, and Interferon-Enriched microglial transcriptional signatures onto the mouse spatial transcriptomic dataset. D) Expression of microglial cell-type transcriptional signatures across spatial tissue domains. E) Comparison of microglial state signature scores between tumor-bearing and sham brains. P-values determined by Wilcoxon rank-sum test. F) Representative spatial feature plots of the PIG signature in tumor-bearing and sham brains. G) Comparison of Early Alzheimer’s Disease (AD)-DAM, Late AD-DAM, and PIG signature scores between tumor-bearing and sham brains. P-values determined by Wilcoxon rank-sum test. PIG, plaque-induced genes; DAM, disease-associated microglia. H) Relative expression of Early AD-DAM, Late AD-DAM, and PIG transcriptional modules across spatial tissue domains.

We next examined the spatial distribution of these microglial programs by mapping signatures comprising the top marker genes of each NMF-defined group onto our spatial transcriptomic dataset (Figure 3C, Table S4). Homeostatic and Early-Activation microglial programs were dispersed throughout tumor and non-tumor domains, whereas the other three programs were almost exclusively expressed within the tumor-associated regions (Figure 3D). However, given the proliferative signature is also expressed by tumor cells, we excluded the Proliferative program from further spatial analysis. When comparing tumor to sham brains, we saw that all microglial programs, except for the Early Activation program, are significantly upregulated in tumor-bearing brains (Figure 3E). Together, these findings reveal spatially organized microglial heterogeneity within the BM microenvironment, characterized by enrichment of interferon-responsive and DAM-like programs in tumor-associated regions.

### Spot deconvolution and cell signature mapping reveal astrocytic and microglial reactivity shared with Alzheimer’s disease models

Recent studies have uncovered pathophysiological links between BM and neurodegenerative disease. We were particularly intrigued by the microglia- and astrocyte-rich cellular architecture observed across our spatial dataset, which is reminiscent of the well-established neuroimmunological response to amyloid beta (Aβ) plaques in Alzheimer’s disease (AD)^17,41^. To explore this connection, we sourced several independently defined AD-associated expression programs and found that they are conserved within our BM samples^42^(Figure 3F-H). Specifically, we find that an Aβ plaque-induced gene signature (PIG), composed of both microglia- and astrocyte-derived inflammatory genes, is elevated in tumor-bearing brains and preferentially localized to tumor-associated spatial domains. Beyond the multicellular PIG signature, we probed for early and late AD-associated microglia DAM programs (AD-DAM), which have previously been described as major tumor cell interactors in BM^31,43^. Of note, these published signatures were evaluated independently of the NMF-derived DAM-like program described above. Both early AD-DAM (*Apoe, B2m, Cstb, Tyrobp, Timp2, H2-D1, Fth1, Lyz2, Ctsb, Ctsd)* and late AD-DAM (*Ank, Spp1, Axl, Csf1, Cst7, Cd9, Cadm1, Clec7a, Ccl6, Itgax, Cd63, Cd68, Ctsa, Lpl, Gusb, Serpine2, Ctsz, Cd52, Ctsl, Hif1a*) programs were upregulated within tumor-bearing brains compared to sham controls and were enriched within tumor-associated spatial domains. Collectively, these results suggest that the neuroinflammatory landscape of BM shares key cellular and molecular hallmarks with the local immune response characteristic of AD models.

### Spatial ligand-receptor inference identifies candidate signaling interactions in the brain metastatic microenvironment

To better understand the extracellular cues shaping tumor-immune interactions within the BM microenvironment, we employed CellChat to infer cell-cell communication events within our spatial transcriptomics dataset^44^. Because individual Visium spots contain multiple cells, this spot-level analysis infers potential ligand-receptor relationships between spot-level spatial regions rather than direct communication between individual cells. Compared to sham brains, the aggregated tumor-bearing dataset contained a greater number and aggregate strength of predicted interactions (Figure S5A). Analysis of relative information flow identified numerous signaling pathways preferentially represented in tumor-bearing brains, while comparison of incoming and outgoing signaling strength revealed distinct predicted communication patterns across intratumoral, interface, and peritumoral domains (Figure S5B-C). Among the most prominent signals in the tumor core were pathways associated with extracellular matrix (ECM) components, including fibronectin (*Fn1*), collagen, and laminin, implicating ECM signaling and possible remodeling as a key feature of the intratumoral niche. We also observed increased predicted amyloid precursor protein (APP) pathway activity in tumor-bearing brains, consistent with our previous findings implicating tumor-associated APP signaling in melanoma BM progression^17^. While similar signaling patterns were detected at the tumor interface and within peritumoral regions, these increasingly extratumoral domains uniquely exhibited enrichment of neurotransmitter-associated pathways, including glutamate and GABA signaling, as well as synaptic regulatory cues such as those driven by neurexins (Figure S5C). Although such signals may simply reflect the presence of nearby neurons, growing evidence suggests that tumor cells can actively engage in neural circuits^45–48^. The presence of inferred neuronal-associated signaling at the BM periphery may therefore create a permissive environment for tumor-neuron crosstalk and structural remodeling, supporting the importance of further investigation into its role in BM progression.

We next examined individual ligand-receptor pairs underlying these predicted domain-level relationships. ECM and cell-contact interactions were robustly detected between intratumoral regions and neighboring intratumoral, interface, and peritumoral spots (Figure S5D-E). These included the Apoe-Trem2/Tyrobp interaction, which has previously been associated with microglial activation, DAM-like transcriptional programs, and phagocytic function^49–51^ (Figure S5D-E). To prioritize candidate immune-related pathways for functional investigation, we focused on ligand-receptor interactions predicted to be enriched in tumor-bearing brains relative to sham controls. This analysis identified several secreted ligands with established associations with macrophage or microglial biology, including *Grn* (granulin), *Spp1* (osteopontin), *Lgals9* (galectin-9), *Psap* (prosaponin) and *Apoe* (apolipoprotein E), with their predicted interactions most prominent in intratumoral and interface domains^52–54^ (Figure 4A). Together, these analyses illustrate a BM microenvironment characterized by spatially organized ECM with robust neuronal- and immune-associated pathways and identify several molecular candidates for subsequent functional investigation.

**Figure 4:**
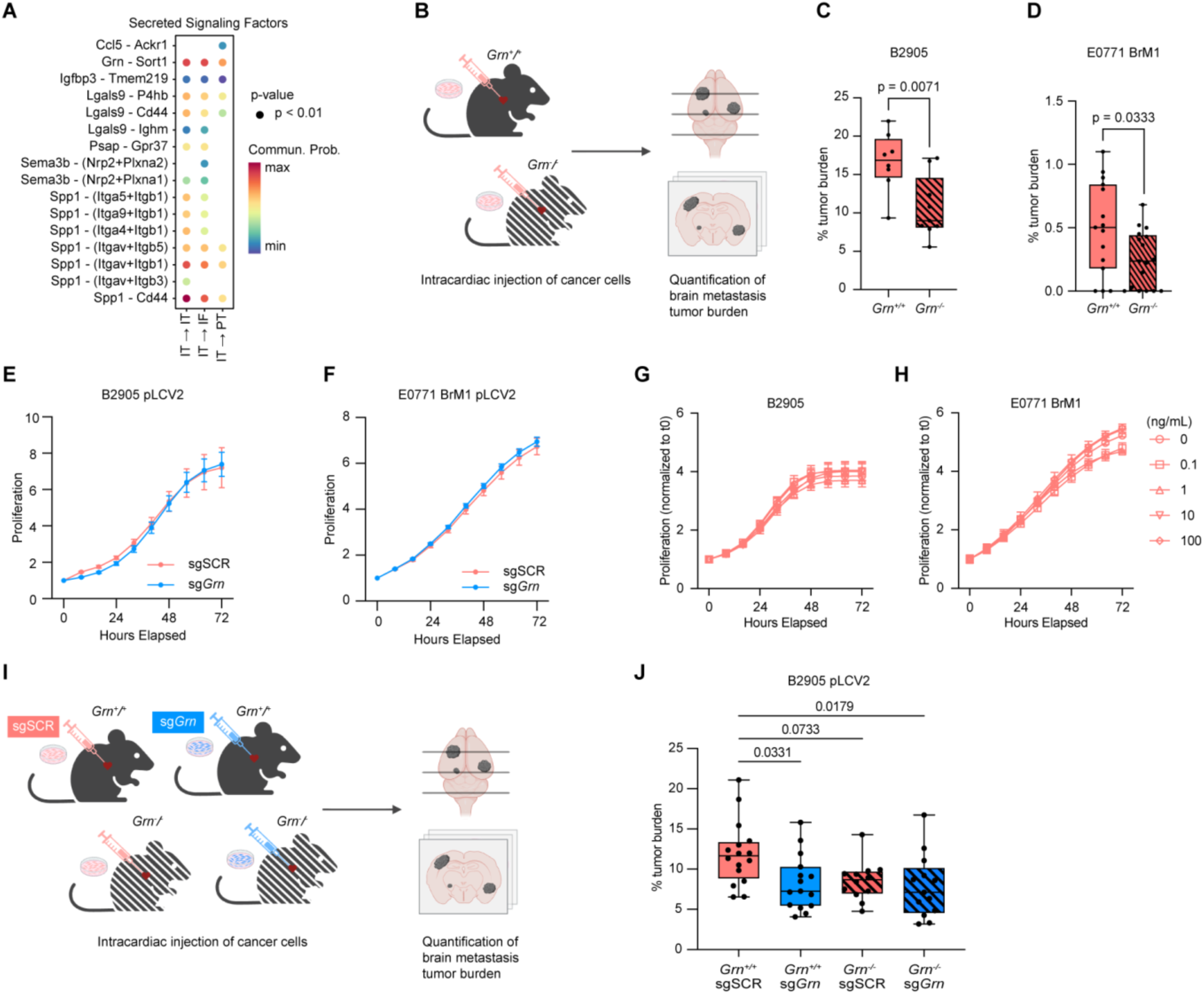
Spatial ligand-receptor analysis identifies progranulin as a candidate regulator of brain metastasis. A) CellChat analysis of predicted enriched ligand-receptor interactions in tumor-bearing brains relative to sham controls across intratumoral (IT), interface (IF), and peritumoral (PT) domains. Dot color indicates predicted communication probability; displayed interactions satisfy p < 0.01. B-D) Experimental schematic for testing the contribution of host-derived PGRN to brain metastasis (B). Tumor cells were injected intracardially into *Grn^+/+^* and *Grn^−/−^* littermates, and brain metastatic burden was quantified at experimental endpoint. Figure generated in BioRender. C-D) Brain metastatic burden in *Grn^+/+^* and *Grn^−/−^* mice following intracardiac injection of B2905 (C) or E0771 BrM1 (D) cells, quantified from H&E-stained sections using HALO image-analysis software. For C-D p-values were calculated using Welch’s two-sided t-test. E-F) In vitro proliferation of pLentiCRISPRv2 (pLCV2) sg*Grn* and control sgSCR B2905 (E) and E0771 BrM1 (F) cells over 72 hours. G-H) Relative proliferation of B2905 (G) and E0771 BrM1 (H) cells following treatment with recombinant mouse PGRN (rmPGRN) in ng/mL. For B2905 (G) no statistically significant differences were observed between groups. For E0771 BrM1 (H) statistically significant differences were observed between groups although not in a biologically meaningful, rPgrn dose-dependent manner. Statistics were calculated by one-way ANOVA with Dunnett’s multiple comparison test comparing each concentration of rmPGRN to 0 ng/mL. I) Experimental schematic evaluating combined host- and tumor-cell *Grn* perturbation. pLCV2 sg*Grn* or pLCV2 sgSCR transduced B2905 tumor cells were injected intracardially into *Grn^+/+^* or *Grn^−/−^* littermates. Figure generated in BioRender. J) Brain metastatic burden quantified from H&E-stained sections using HALO image-analysis software. Statistical comparisons were performed using one-way ANOVA with Tukey’s multiple comparisons test.

### Progranulin loss results in reduced brain metastasis

Among the candidate immune-associated pathways identified by our spatial analyses, we prioritized progranulin (PGRN), a secreted 88-kDa glycoprotein encoded by *Grn*. *Grn* is a component of the plaque-induced gene signature enriched in the tumor-bearing samples, and PGRN has established roles in microglial inflammatory and lysosomal biology. In humans, heterozygous loss-of-function variants in *GRN* cause a familial form of frontotemporal dementia, while biallelic loss-of-function causes neuronal ceroid lipofuscinosis, a lysosomal storage disease^55–57^. These observations have motivated therapeutic strategies to restore PGRN availability in neurodegenerative disease, including targeting sortilin (SORT1), a receptor that mediates PGRN internalization and lysosomal trafficking, to reduce PGRN clearance and increase extracellular PGRN levels^58,59^.

Given the association between PGRN and microglial inflammatory activity, we hypothesized that targeting PGRN may influence BM progression by attenuating immunosuppressive signaling and promoting a more immunostimulatory, anti-tumor landscape. To test this hypothesis, we injected our brain-tropic tumor cell lines intracardially into *Grn^−/−^* mice and *Grn^+/+^*littermate controls and measured BM burden at a humane experimental endpoint (Figure 4B). Following injection of B2905 melanoma cells, BM burden was significantly reduced in *Grn^−/−^* hosts compared with wild-type littermates (p = 0.0071; Figure 4C). E0771 BrM1 breast cancer cells exhibited similarly reduced brain metastatic capacity in *Grn^−/−^*hosts (p = 0.0333; Figure 4D). These findings suggest that host-derived PGRN may play a pro-tumorigenic role in the BM microenvironment (Figure 4C-D).

Importantly, multiple groups have identified protumoral functions of tumor cell-derived PGRN in other cancer models^60–63^. To investigate whether tumor-derived PGRN directly influences tumor cell growth in our BM models, we generated *Grn*-knockout B2905 and E0771 BrM1 cells using CRISPR/Cas9. CRISPR-mediated knockout of *Grn* achieved significant reduction in PGRN levels, with no significant differences in in vitro proliferation compared to control cells (Figures 4E-F, S6A-B). Conversely, to mimic the availability of microenvironmental PGRN, we treated cells with recombinant murine progranulin (rmPGRN) and observed no dose-dependent change in proliferation in vitro (Figure 4G-H).

While tumor-derived PGRN does not play an intrinsically mitogenic role in our BM models in vitro, secreted tumor-derived PGRN may still actively shape the BM microenvironment. To simultaneously investigate the contribution of host- and tumor-derived PGRN in vivo, we injected *Grn* knockout and control B2905 melanoma tumor cells into *Grn*^−/−^ mice and *Grn*^+/+^ littermate controls (Figure 4I). In this cohort, tumor-cell *Grn* loss produced a significant reduction in BM burden among *Grn*^+/+^ mice (p = 0.0331), and combined host- and tumor-cell *Grn* loss also significantly reduced BM burden relative to the fully *Grn*-proficient condition (p = 0.0179; Figure 4J). Together, our findings support the possibility of a joint role for both tumor-derived and host-derived PGRN in BM progression, and motivate further investigation into the mechanism through which stromal- and tumor-derived PGRN may shape the BM tumor microenvironment.

### GRN expression is associated with a conserved lysosomal and DAM-like transcriptional program in mouse and human brain metastasis

To define the cellular and spatial context of *Grn* expression in BM, we first examined its distribution across our murine spatial transcriptomic atlas. Across all models *Grn* expression was more prominent in tumor-bearing brains than in sham-injected controls and was upregulated across all tumor-associated tissue domains, likely reflecting contributions from both tumor cells and changes in the surrounding myeloid compartment (Figures 5A, S7A). In the scRNA-seq reference data, *Grn* expression was enriched in myeloid populations, particularly in microglia^27^ (Figure 5B). Among the NMF-defined microglial groups, *Grn* expression was most enriched in DAM-like microglia, followed by interferon-enriched microglia, associating *Grn* with transcriptionally reprogrammed microglial states in the BM microenvironment (Figure 5C).

**Figure 5:**
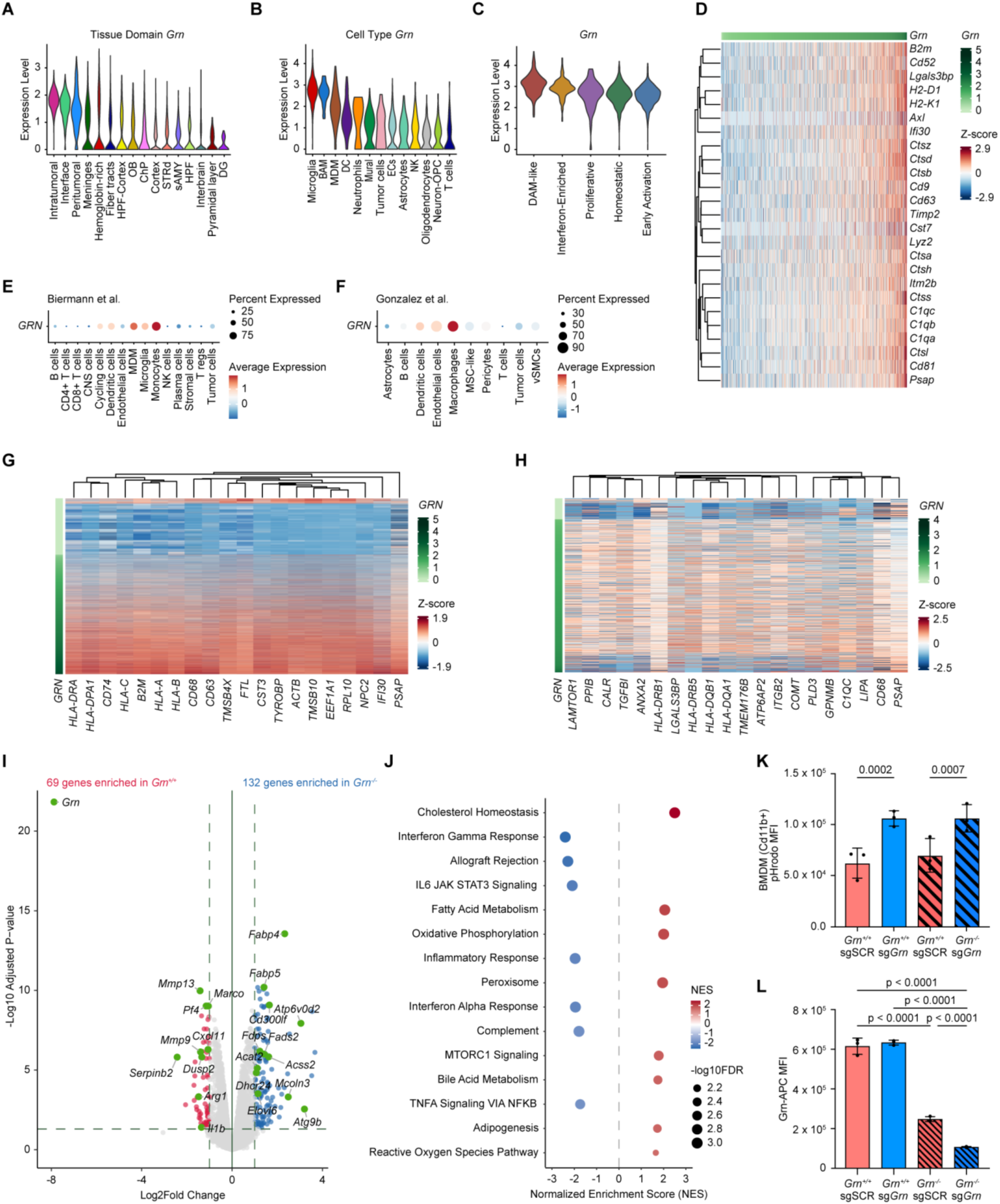
*GRN* expression is associated with conserved antigen-presentation, lysosomal, and DAM-like myeloid programs in mouse and human brain metastases. A) *Grn* expression across annotated spatial tissue domains in the mouse brain metastasis spatial transcriptomics atlas. B) *Grn* expression across tumor, immune, and stromal cell types, using Rodriguez-Baena et al., 2025 reference scRNA-seq data^27^. BAM, border-associated macrophage; MDM, monocyte-derived macrophage; DAM, disease-associated microglia; DC, dendritic cell; ECs, endothelial cell; OPC, oligodendrocyte precursor cell. C) *Grn* expression across the five NMF-defined microglial states identified in the Rodriguez-Baena et al. mouse brain metastasis scRNA-seq dataset. D) Heatmap showing scaled expression of top 25 genes most positively correlated with *Grn* in mouse brain metastasis-associated microglia. E-F) Dot plots showing *GRN* expression across annotated cell populations in the human melanoma brain metastasis scRNA-seq dataset from (E) Biermann et al., and the multi-tumor brain metastasis dataset from (F) Gonzalez et al^7,9^. G-H) Heatmaps showing scaled expression of top 20 *GRN*-correlated genes within myeloid populations from the Biermann et al. (G) and Gonzalez et al. (H) datasets. Cells are ordered according to *GRN* expression^7,9^. I) Volcano plot showing differential gene expression between *Grn*^+/+^ and *Grn*^−/−^ bone marrow-derived macrophages (BMDMs) assessed by bulk RNA sequencing. Genes significantly enriched in *Grn*^+/+^ (red) or *Grn*^−/−^ (blue) BMDMs are highlighted; dashed lines denote the applied fold-change and adjusted p-value thresholds. J) Hallmark gene-set enrichment analysis comparing *Grn*^−/−^ and *Grn*^+/+^ BMDMs. Positive normalized enrichment scores indicate enrichment in *Grn*^−/−^ BMDMs, negative scores indicate enrichment in *Grn*^+/+^ BMDMs. K) Detection of pHrodo-labeled control pLentiCRISPRv2 (pLCV2) sgSCR or *Grn*-deficient sg*Grn* B2905 tumor cells within *Grn*^+/+^ and *Grn*^−/−^ BMDMs, quantified by flow cytometry as pHrodo mean fluorescence intensity (MFI) within CD11b+ BMDMs. Individual points represent technical replicates; bars show mean ± SD. Statistical comparisons were performed using one-way ANOVA and Tukey’s multiple comparison test. L) Intracellular PGRN abundance in CD11b+ BMDMs following coculture with control pLCV2 sgSCR or sg*Grn* B2905 cells, quantified by flow cytometry as APC MFI. Individual points represent technical replicate measurements; bars show mean ± SD. Statistical comparisons were performed using one-way ANOVA and Tukey’s multiple comparison test.

To further resolve the transcriptional programs associated with *Grn*, we identified genes whose expression correlated most strongly with *Grn* within microglia. Notably, both early-stage AD-DAM genes (*B2m, Timp2, H2-D1, Lyz2, Ctsb, Ctsd*) and late-stage AD-DAM genes (*Axl, Cst7, Cd9, Cd63, Ctsa, Ctsz, Ctsl, Cd52*) were among the top correlates^31^ (Figure 5D). *Grn* expression also correlated with key genes involved in antigen processing and presentation, including its binding partner prosaposin (*Psap*), which facilitates shuttling of PGRN to the cell surface and endosomes, and MHC molecules (*B2m, H2-D1, H2-K1*)^55^. Additionally, *Grn* showed strong associations with genes encoding lysosomal enzymes (*Ifi30, Ctsh, Ctss, Ctsl, Ctsa, Ctsz, Ctsd, Ctsb*), components of the C1 complement complex (*C1qa, C1qb, C1qc*), and exosomal markers (*Cd9, Cd81*)^64^ (Figure 5D). Together, these findings associate *Grn* expression with a DAM-like transcriptional program encompassing lysosomal function, antigen processing and presentation, and complement signaling.

To better understand the potential mechanisms through which PGRN participates in intercellular communication within the BM microenvironment, we employed NICHES and LIANA, two complementary tools for inferring ligand-receptor interactions in our spatial transcriptomics dataset^65,66^. Our analyses revealed colocalization of *Grn-Sort1* and *Grn-Tnfrsf1a* within intratumoral regions, implicating both *Sort1* (sortilin) and *Tnfrsf1a* (TNF receptor 1) as potential mediators of PGRN-associated signaling in BM (Figure S7B-C). Consistent with our gene correlation analyses, these findings suggest that PGRN may exert its effects by modulating endolysosomal trafficking via Sort1 and inflammatory signaling via Tnfrsf1a^67,68^.

To determine whether *GRN*-associated myeloid programs were also present in human BM, we analyzed two independent scRNA-seq datasets spanning multiple primary tumor types. The first dataset, published by Biermann et al., profiled 22 treatment-naïve melanoma BMs (MBM) and 10 extracranial melanoma peripheral metastases (MPM)^7^. In MBM samples, *GRN* expression was predominantly enriched in the myeloid compartment, particularly within monocytes and MDMs (Figure 5E). The second dataset, from Gonzalez et al., profiled 15 human parenchymal BMs from multiple primary tumor types with *GRN* again robustly expressed in macrophages^9^ (Figure 5F). In their original report, Gonzalez et al. identified two transcriptionally distinct subsets of metastasis-associated macrophages (MAMs): APOE^+^ MAMs, which expressed an immunomodulatory signature including C1Q complement chains, *SPP1*, and HLA-related genes; and S100A8^+^ MAMs, characterized by a pro-inflammatory gene program, including *S100A8/9*, *CXCL8*, and *FCN1*, with low HLA expression. Notably, *GRN* expression was higher in the APOE^+^ MAMs, consistent with its association with APOE, C1Q, and HLA-related programs (Figure S8A).

Finally, we examined genes correlated with *GRN* expression in the macrophage and microglial compartments of both human datasets. Across datasets, *GRN* expression correlated with genes associated with macrophage identity and lysosomal function, including *CD68*, *PSAP*, *LIPA*, *PLD3*, and cathepsins such as *CTSB* and *CTSD*. Additional correlates included HLA class I and II genes, *B2M*, *APOE*, and C1q complement components *C1QA*, *C1QB*, and *C1QC* (Figure 5G-H). This transcriptional profile closely resembled the DAM-like program observed in our mouse models. Functional enrichment among *GRN*-correlated genes identified pathways related to phagocytosis, lysosomal function, antigen processing and presentation, and oxidative phosphorylation (Figure S8B-C). Together, these analyses support a conserved association between *GRN* expression and lysosomal, antigen-processing, complement-associated, and other features of DAM-like myeloid programs in mouse and human BM.

### Loss of progranulin alters macrophage metabolic and phagocytic programs

Our in vivo experiments implicated both host- and tumor-derived PGRN in BM progression, while our cross-species computational analyses associated *GRN* expression with lysosomal and DAM-like myeloid programs. Building on these findings, we next sought to determine whether *Grn* loss alters macrophage phenotype and function in vitro. We used bone marrow-derived macrophages (BMDMs) cultured from non-tumor-bearing *Grn*^−/−^ mice and *Grn*^+/+^ littermates and performed bulk RNA sequencing. Differential-expression analysis identified 132 genes upregulated and 69 genes downregulated in *Grn*^−/−^ BMDMs relative to *Grn*^+/+^ BMDMs (Figure 5I).

Gene-set enrichment analysis revealed enrichment of oxidative phosphorylation, cholesterol homeostasis, fatty-acid metabolism, and peroxisomal programs in *Grn*^−/−^ BMDMs. Conversely, *Grn*^+/+^ BMDMs were enriched for inflammatory programs, including interferon-α and interferon-γ responses, complement, and TNF-α signaling through NF-κB (Figure 5J). Thus, *Grn*-deficient BMDMs showed relative enrichment of oxidative and lipid-metabolic gene sets, whereas wild-type BMDMs showed enrichment of inflammatory gene sets. The enrichment of interferon-response and complement programs in wild-type BMDMs was directionally consistent with their positive association with *GRN* expression in our prior mouse and human analyses.

Having characterized the transcriptional effects of *Grn* loss in BMDMs, we next examined its effects on tumor-macrophage interactions in vitro. Because PGRN regulates endolysosomal biology, and microglial phagocytosis has been implicated in restricting early BM colonization, we examined whether *Grn* status affects macrophage engulfment of tumor cells^69^. We used a pHrodo-based assay to quantify tumor-derived material within acidic compartments of *Grn*^+/+^ and *Grn*^−/−^ BMDMs following coculture with sgSCR control or sg*Grn* B2905 tumor cells. When cocultured with sgSCR tumor cells, *Grn^+/+^* and *Grn^−/−^* BMDMs exhibited similar pHrodo fluorescence. In contrast, tumor-cell *Grn* loss significantly increased pHrodo signal in BMDMs of either genotype (Figures 5K, S9A). These findings associate tumor-cell *Grn* loss with increased pHrodo signal in macrophages, which may reflect increased uptake, altered acidification or phagolysosomal processing (due to prolonged persistence of labeled tumor-derived material), or a combination of these effects.

In parallel, flow cytometric analysis of *Grn^−/−^* and *Grn^+/+^* BMDMs co-cultured with sgSCR or sg*Grn* tumor cells demonstrated that intracellular PGRN levels within macrophages are dependent on both macrophage and tumor cell expression of *Grn* (Figures 5L, S9B). This observation is consistent with the possibility that tumor-derived PGRN may contribute to the intracellular PGRN pool within macrophages. Ex vivo analysis of brain metastases similarly showed that microglial PGRN signal was lowest in *Grn*−/− hosts bearing sg*Grn* tumors, supporting contributions from both host and tumor-derived PGRN within the BM microenvironment (Figures S9C-D). Overall, these findings suggest that both macrophage- and tumor-derived PGRN may influence tumor cell engulfment, with tumor-cell *Grn* status exerting a particularly prominent effect in vitro.

Together, these data associate *Grn* loss with altered macrophage metabolic programs and show that tumor-cell *Grn* impairs the accumulation and processing of tumor-derived material by macrophages in vitro. They further suggest that tumor-derived PGRN may contribute to the PGRN pool detected within neighboring myeloid cells. Although these experiments do not establish that increased tumor-cell phagocytosis accounts for the reduced BM burden observed in vivo, they nominate macrophage phagolysosomal function as a candidate mechanism connecting PGRN to tumor-myeloid interactions and increased BM burden.

### GRN-high myeloid niches colocalize with immune suppressive programs and dysfunctional CD8+ T cell states in human brain metastases

In light of the transcriptomic association between *GRN* and immunomodulatory myeloid states in our mouse spatial transcriptomic atlas and analysis of human scRNA-seq data, we next asked whether *GRN*-high myeloid populations occupy spatially distinct immune niches in human BM.

To explore this question, we analyzed 10x Visium spatial transcriptomic data from 16 human melanoma BM samples representing 13 patients. Within each sample, we identified spots enriched for microglial or MDM transcriptional signatures, using human orthologs of the gene signatures derived from the mouse scRNA-seq dataset^27^ (Table S5). Myeloid-enriched spots were stratified as *GRN*-high and *GRN*-low groups based on the upper and lower 20th percentiles of *GRN* expression, respectively (Figure S10A). We then examined the spatial relationship of *GRN*-high and -low spots to neighboring spots by applying transcriptional signature scores for immunosuppression and antigen presentation (Table S6). *GRN*-high microglia- and MDM-enriched spots were frequently localized within neighborhoods with high immunosuppression and antigen presentation scores. Across samples, *GRN*-high myeloid spots had more immediate neighboring spots with high immunosuppression scores than their *GRN*-low counterparts, a relationship observed for both microglia- and MDM-enriched spots (Figures 6A-C, S10B). Similarly, *GRN*-high microglia- and MDM-enriched spots exhibited significantly greater spatial association with antigen presentation-high neighboring spots compared with *GRN*- low myeloid spots (Figure 6B-C). Together, these findings demonstrate that elevated *GRN* expression within the myeloid compartment is associated with spatially organized immune-regulatory niches characterized by both immunosuppressive and antigen-presentation programs in human BM. Because these programs can shape T cell activation and exhaustion, we next used single-cell-resolution spatial profiling to determine whether *GRN*-high myeloid populations were also associated with dysfunctional T cell states.

**Figure 6:**
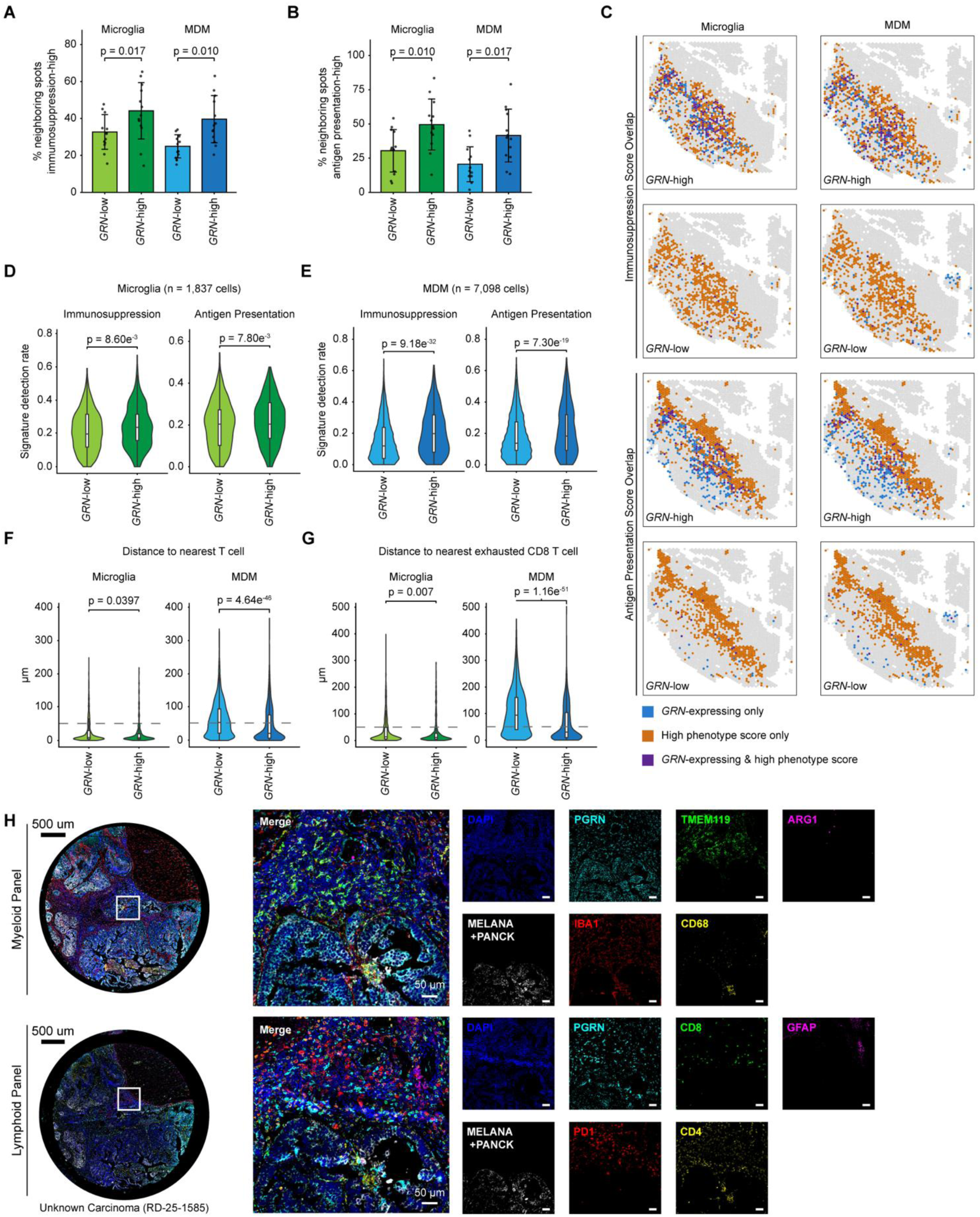
*GRN*-high myeloid niches are spatially associated with immunosuppressive and dysfunctional T cell states in human brain metastases. A-B) Analysis of human melanoma brain metastasis Visium data from 16 samples representing 13 patients. Percentage of immediate neighboring spots with high immunosuppression (A) or antigen-presentation (B) signature scores surrounding *GRN*-low and *GRN*-high microglia- or monocyte-derived macrophage (MDM)-enriched spots. Each point represents an individual patient; bars show mean ± SD. P-values calculated using Wilcoxon signed-rank test. C) Representative spatial maps showing the overlap between *GRN*-low or *GRN*-high microglia- and MDM-enriched spots and spots with high immunosuppression or antigen-presentation signature scores. Blue indicates *GRN*-expressing myeloid spots only, orange indicates spots with high signature scores only, and purple indicates spatial overlap between the two classifications. D-E) Violin plots comparing anti-inflammatory and antigen-presentation signature detection rates between *GRN*-low and *GRN*-high microglia (D; n = 1,837 cells) and macrophages (E; n = 7,098 cells) in Xenium spatial transcriptomic data from three human brain metastases. P-values calculated using BH-adjusted one-sided Wilcoxon test. F-G) Distance from *GRN*-low and *GRN*-high microglia or macrophages to the nearest T cell (F) or exhausted CD8+ T cell (G) in the Xenium dataset. Dashed lines indicate 50-µm neighborhood distances. P-values calculated using BH-adjusted one-sided Wilcoxon test. H) Multiplex immunofluorescence staining of the human brain metastasis RD-25-1585 (unknown carcinoma primary) analyzed using Xenium spatial transcriptomics. Scale bars are 50µm unless otherwise indicated.

To achieve single-cell spatial resolution, we analyzed Xenium spatial transcriptomic data collected from three human BM originating from melanoma (n=1), breast cancer (n=1) and a carcinoma of unknown primary origin (n=1). Consistent with our previous analyses, *GRN* expression was enriched within MDM and microglia populations (Figure S11A). Furthermore, we again observed that *GRN*-high MDM and microglia were more frequently proximal to cells that scored highly for immunosuppression and antigen presentation programs (Figures 6D-E, S11B). Direct proximity analyses further showed that microglia were globally located closer to T cells than MDMs (Figure 6F). Furthermore, among both microglia and MDM, *GRN*-high cells were more frequently found near CD8+ T cells annotated as exhausted (PD-1+, LAG3+, TIM-3+, CTLA4+, TIGIT+, ENTPD1+, TOX+, NR4A1+) when compared to their *GRN*-low counterparts (Figure 6G). These findings suggest that *GRN*-rich cellular neighborhoods are predominantly immunosuppressed and associated with exhausted T cell states within human BM.

We next used multiplex immunofluorescence to investigate the spatial relationship between dysfunctional T cell phenotypes and *GRN*-expressing microglia and MDMs at the protein level in the same human BM specimens analyzed by Xenium. IBA1+TMEM119+ microglia and IBA1+TMEM119-macrophages universally stained positive for PGRN, as did MELANA/PANCK+ tumors. Representative regions highlight PD-1+ CD8+ T cells within or adjacent to regions containing IBA1+TMEM119+ microglia and IBA1+TMEM119-macrophages (Figure 6H). This is consistent with the finding that *GRN*-high myeloid cells occupy neighborhoods enriched for dysfunctional PD1+ CD8+ T cell states. Together, the human spatial transcriptomic analyses, supported by protein-level observations, associate *GRN*-rich myeloid niches with immunoregulatory programs and dysfunctional T cell states in human BM.

## Discussion

In this study, we established a panel of syngeneic preclinical models to characterize the spatial transcriptomic landscape of BM, ultimately identifying PGRN as a novel candidate therapeutic target for BM. Many studies have employed immunocompromised models to explore tumor cell-intrinsic adaptations to the BM microenvironment. Yet, the implementation of ICIs as first-line treatment for BM of various tumor origins underscores the importance of exploring tumor cell-extrinsic adaptations of the microenvironment. Using our syngeneic preclinical models, we describe spatially localized myeloid and astrocytic responses, with interferon-enriched and DAM-like programs concentrated in the immediate tumor microenvironment. The prominence of microglia and MDMs in our models is consistent with studies identifying myeloid cells as a major component of human BM^7,9,11,12^. Preclinical models have further supported the functional significance of these populations, as genetic and pharmacologic depletion of tumor-associated microglia has repeatedly been shown to reduce tumor burden^13,27,70^. Our atlas adds spatial context by showing that interferon-enriched and DAM-like myeloid programs are differentially distributed across tumor-associated regions. These observations complement studies implicating *Rela*/NF-κB signaling and tumor-derived tenascin-C in the emergence of inflammatory and DAM-like microglial states in BM mouse models^27,43^.

PGRN provides a potential molecular connection between these myeloid programs and brain metastatic progression. PGRN regulates lysosomal homeostasis in the central nervous system. Inherited *GRN* loss of function causes neurodegenerative disease accompanied by lysosomal dysfunction and altered glial activation, ultimately driving neuronal toxicity and proteinopathies^55,71,72^. Given the link between PGRN deficiency and heightened neuroinflammation, we hypothesized that reducing PGRN availability could enhance anti-tumor immune responses and offer therapeutic benefit in the context of BM. Prior studies have connected PGRN to macrophage efferocytosis, fibrotic remodeling, and cytotoxic T cell exclusion in other cancer types, although the relative contribution of tumor- and host-derived PGRN varies by tumor context across independent studies^73–76^.

In our study, we found that host- and tumor-cell *Grn* loss both contribute to reduced BM burden, supporting a net tumor-promoting role for PGRN in BM. Across mouse and human datasets, *GRN* expression was associated with DAM-like transcriptional states as well as with genes involved in lysosomal function, phagocytosis, complement signaling, and antigen processing and presentation. Interestingly, in malignant brain tumors, tumor-associated microglia and MDM antigen-presentation machinery can be upregulated but not functionally active because of co-induced immunosuppressive signaling^8,77^. Persistent antigen presentation within this context has been shown to contribute to CD8+ T cell dysfunction^78^, whereas blockade of the anti-phagocytic checkpoints CD47 and CD24 can restore CD8+ T cell reactivity via cross-presentation of tumor antigens^79^. This framework linking immunosuppressive tumor-associated microglia/MDMs with attenuated T cell-mediated adaptive immunity in the brain tumor microenvironment aligns with our findings regarding the proximity of *GRN*-high microglia and MDM populations, associated with elevated antigen presentation signatures, to dysfunctional CD8+ T cell states in human BM.

Our in vivo and in vitro functional studies begin to shed light on how PGRN may influence macrophage phenotypes in BM. In vitro, *Grn*-deficient BMDMs exhibited altered metabolic and lipid-homeostatic programs, and tumor-cell *Grn* loss led to increased detection of pHrodo-labeled tumor-derived material within acidic compartments of BMDMs of either genotype. Of note, increased detection of pHrodo-labeled tumor-derived material within acidic compartments of BMDMs may be plausibly explained by multiple mechanisms, including increased capacity for tumor engulfment, altered phagolysosomal processing, or aberrant persistence of labeled cargo^80–82^. This phenotype could therefore reflect altered susceptibility of *Grn*-deficient tumor cells to macrophage engulfment, a cell-extrinsic effect of tumor-derived PGRN on macrophage function, altered macrophage processing of tumor-derived material, or a combination of these mechanisms. Previous studies have shown that PGRN deficiency can enhance macrophage phagocytosis or efferocytosis while impairing lysosomal acidification, autophagy, or degradative function^80–82^. Together with the association between GRN and antigen-processing and presentation programs, these observations raise the possibility that PGRN influences antigen handling by regulating phagocytic uptake or subsequent phagolysosomal processing.

The human spatial analyses further extend our findings by placing *GRN*-associated myeloid biology within an adaptive immune context in BM. In human BM, *GRN*-high myeloid-enriched regions were more closely associated with transcriptional signatures of immune suppression and antigen presentation than *GRN*-low regions. Higher-resolution spatial profiling additionally placed *GRN*-high macrophages near CD8+ T cell states associated with exhaustion. The recurrence of this phenomenon across independent spatial platforms and patient cohorts suggests that *GRN*-high myeloid populations occupy distinct immune neighborhoods that may contribute to local regulation of T cell activity in BM.

Multiple independent groups have now described pro-tumorigenic roles for PGRN both in vitro and in vivo, culminating in the development of PGRN-targeting therapies for clinical use^61,62^. Currently, an anti-PGRN/GP88 monoclonal antibody, AG01, has entered Phase 1 evaluation in patients with advanced solid tumors^63^. Beyond our studies, additional work is warranted to further determine whether pharmacologic PGRN blockade can demonstrate therapeutic benefit in BM, either alone or in addition to immune checkpoint inhibitors. Our nomination of candidate *Grn-Sort1* interactions also suggests SORT1 as an additional, context-dependent therapeutic entry point. In fact, SORT1-directed antibodies reached Phase 2-3 clinical testing in neurodegenerative disease, though they demonstrated a lack of efficacy for those indications^83,84^; future studies in the specific context of BM are needed to determine the anti-tumor efficacy of SORT1 modulation.

Collectively, this study provides a cross-tumor immunocompetent model panel and spatial transcriptomic atlas for identifying shared features of the BM microenvironment. We are confident that our development of a panel of syngeneic BM models and subsequent querying of the BM microenvironment via spatial transcriptomics will be a valuable resource to the field. We further extend this resource through complementary single-cell, Visium, Xenium, and multiplex immunofluorescence analyses of human BM, providing a multi-platform view of conserved myeloid programs and their spatial relationships with PGRN expression and adaptive immune states. The convergence of mouse perturbation studies with independent human single-cell spatial transcriptomic analyses supports PGRN as a candidate therapeutic vulnerability in BM. Future mechanistic studies defining how tumor- and host-derived PGRN influence myeloid function, and deeper exploration of the mechanistic link to adaptive immune responses, may reveal new opportunities for therapeutic intervention within the BM immune niche.

---

**Figure S1:**
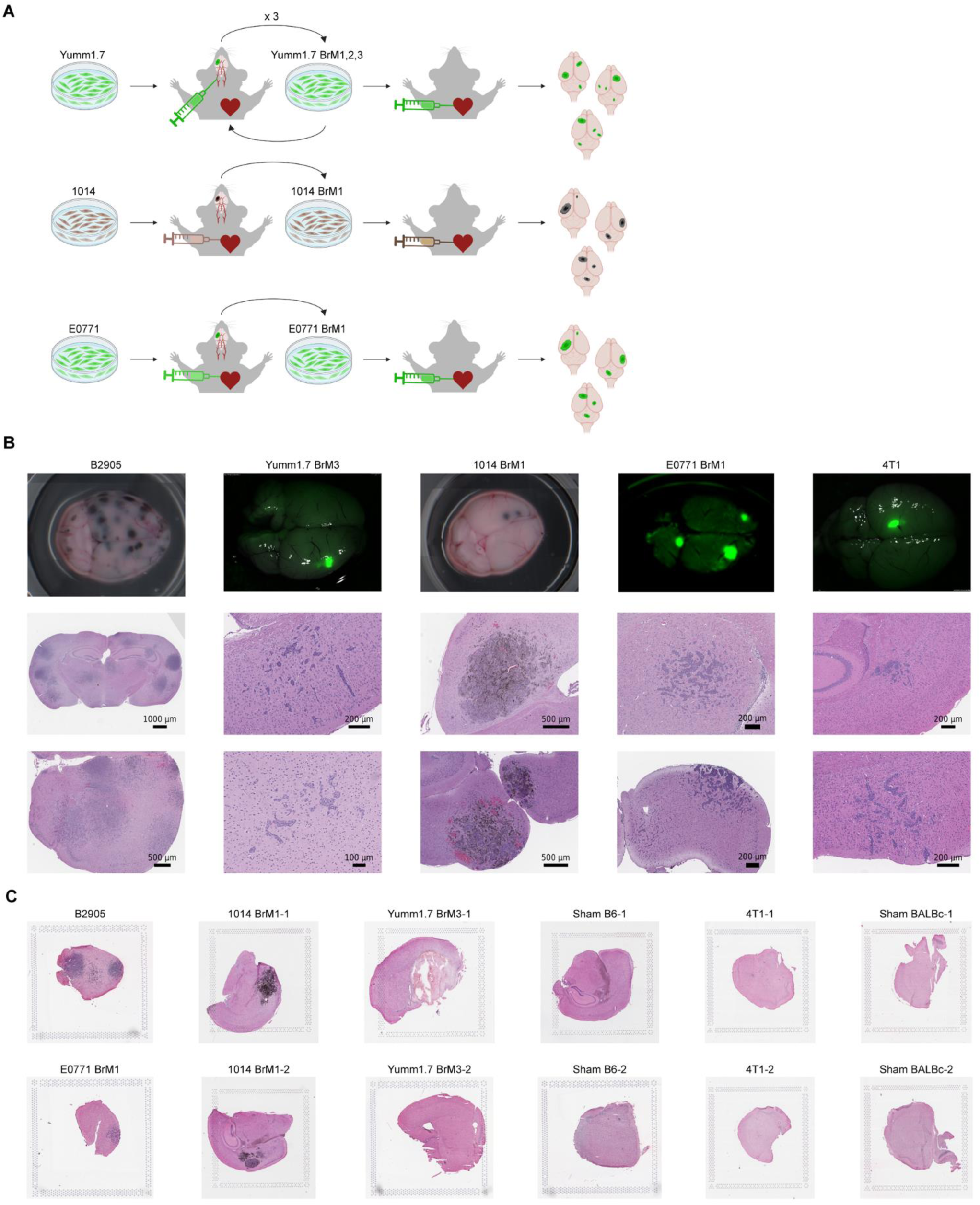
Generation of a syngeneic mouse brain metastasis model panel for spatial transcriptomic analysis. A) Generation of brain-tropic cell lines. Yumm1.7 BrM3 was derived from Yumm1.7 cells stably transduced with a GFP-luciferase reporter. Yumm1.7 GFP+ cells were injected into mice via the internal carotid artery. Parenchymal brain metastases were harvested from mice, expanded in vitro, and subjected to two subsequent rounds of serial intracardiac injection. After the third round of in vivo passage, harvested GFP+ cells were expanded in vitro as Yumm1.7 BrM3 cells. For E0771 cells (also transduced to express GFP-luciferase) and 1014 cells, only one round of serial injection was needed to yield highly brain-tropic lines. Here, the cells were injected intracardially, and the resulting brain metastases were harvested and expanded in culture to yield E0771 BrM1 and 1014 BrM1 cells. Figure generated in BioRender. B) Representative gross and histologic appearance of brain metastases generated by the five syngeneic models. Gross lesions were visualized by pigmentation in melanocytic models or GFP fluorescence in GFP-labeled models. C) H&E of samples used for Visium spatial transcriptomic sequencing and analysis.

**Figure S2:**
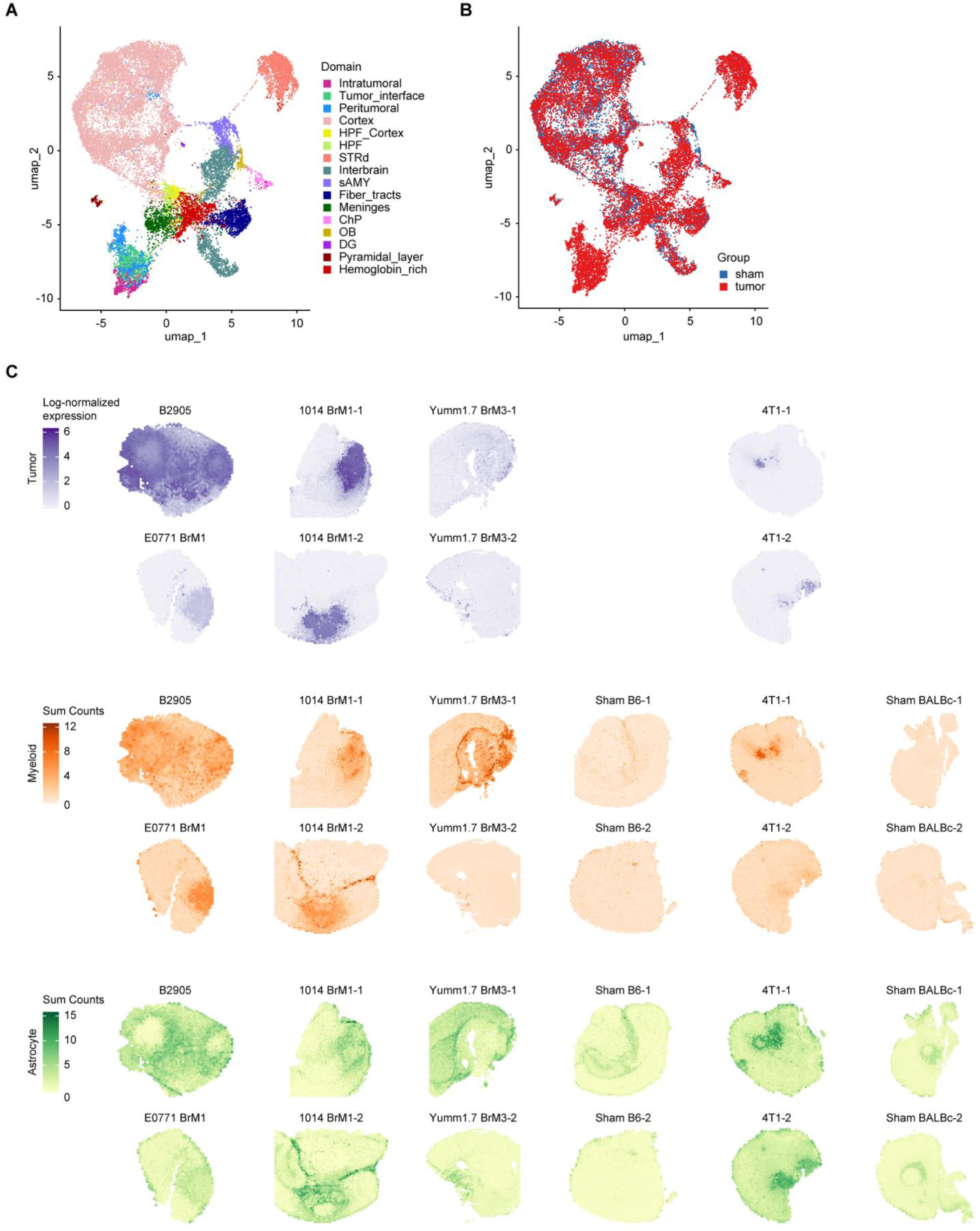
Marker expression supports spatial transcriptomic tissue-domain annotation. A-B) UMAP projection of spatial transcriptomic spots, colored by tissue-domain assignment (A) or experimental group (B) tumor-bearing versus sham. C) Spatial expression of tumor-lineage markers and myeloid and astrocytic transcriptional signatures across tumor-bearing and sham samples. Tumor markers were *Melana* for B2905 and 1014 BrM1, *Mgp* for Yumm1.7 BrM3, and *Krt8* for 4T1 and E0771 BrM1. The myeloid score comprised *Aif1, Cd68, Trem2,* and *Lyz2*, and the astrocyte score comprised *Gfap, Aqp4,* and *Serpina3n*.

**Figure S3:**
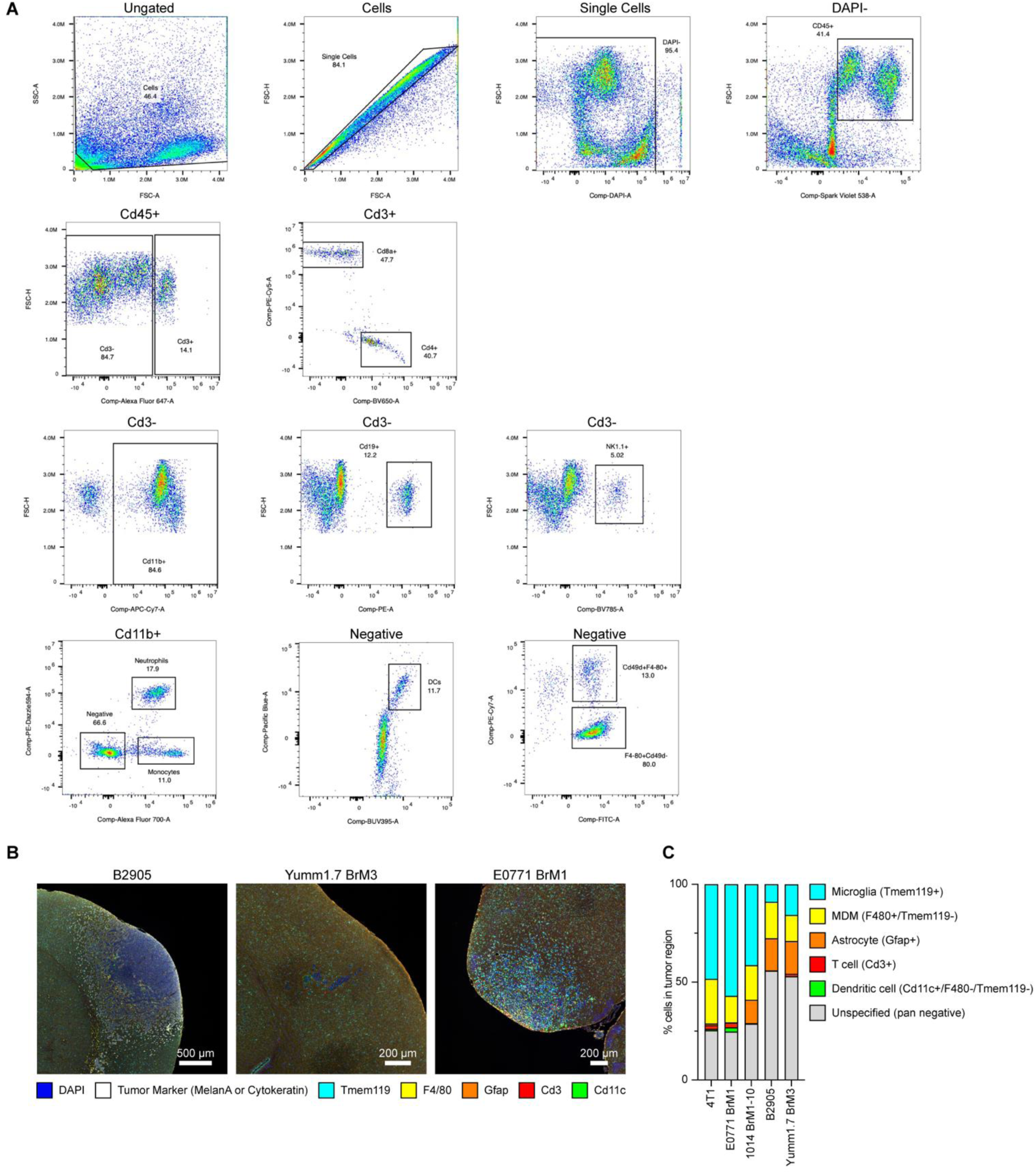
Orthogonal immune profiling supports a myeloid-rich brain metastatic microenvironment. A) Flow cytometry gating strategy used to identify major immune cell populations within whole mouse cerebrums bearing B2905-derived brain metastases. B) Additional representative multiplex IF staining and cell-type quantification across E0771 BrM1, Yumm1.7 BrM3, and B2905 brain metastasis models. TMEM119, F4/80, and CD11c mark myeloid populations; CD3 marks T cells; GFAP marks reactive astrocytes; and DAPI identifies nuclei. Tumor cells are identified using cytokeratin or Melana as appropriate for tumor lineage. Scale bars, 200 μm. “Unspecified” denotes DAPI+ cells negative for the assessed lineage markers. C) Quantification of cell populations within tumor regions based on IF staining. “Unspecified” denotes DAPI+ cells that did not stain positively for the assessed cell identity markers and may include tumor, immune, or non-immune stromal cells.

**Figure S4:**
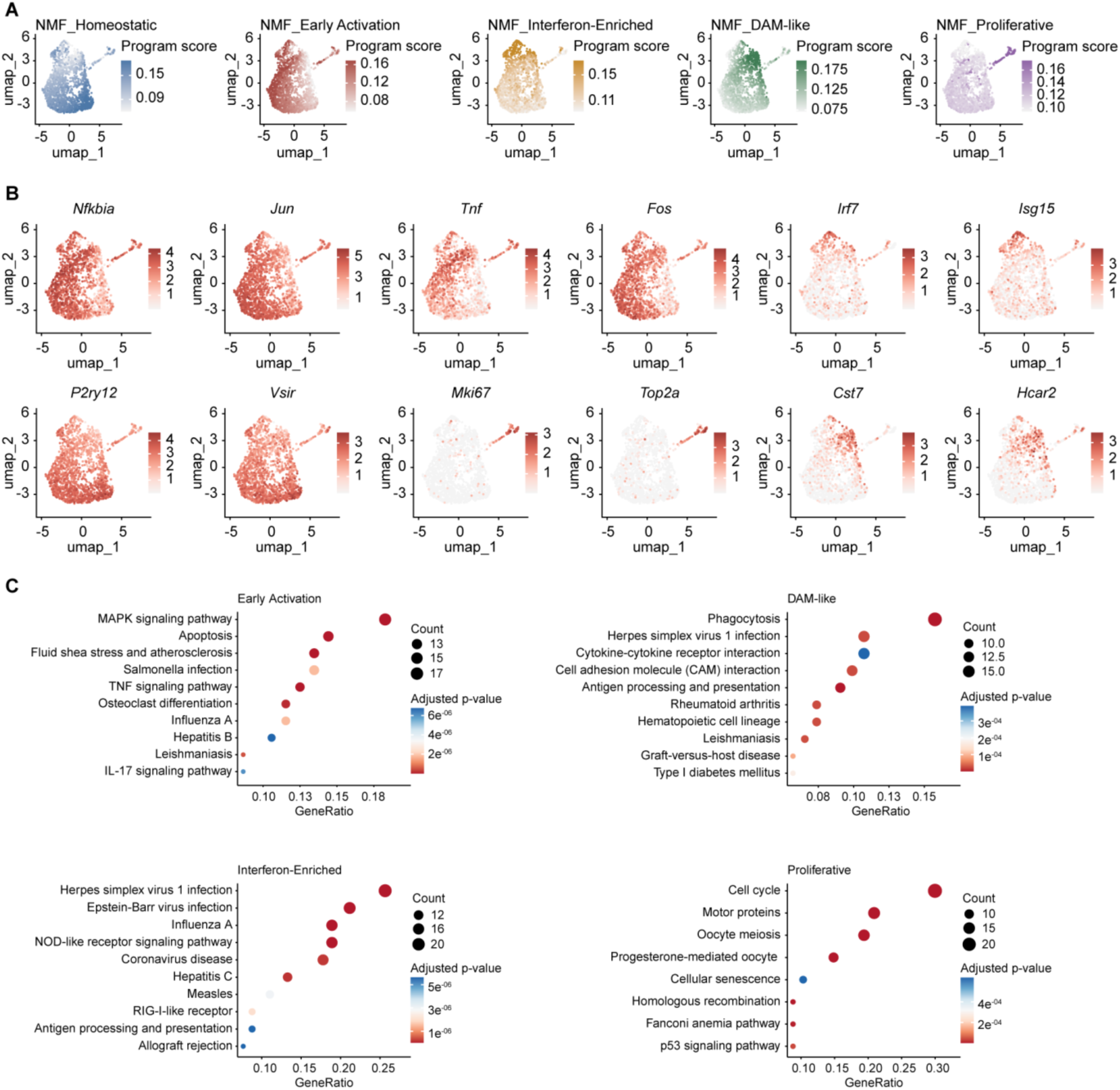
NMF resolves transcriptionally distinct microglial states in brain metastasis. A) UMAP feature plots showing per-cell scores for the five NMF-derived microglial transcriptional programs: Homeostatic, Early Activation, Interferon-Enriched, DAM-like, and Proliferative. B) UMAP feature plots showing expression of representative genes associated with the identified microglial states. C) KEGG pathway enrichment analysis of the top 200 differentially expressed genes from each NMF-defined microglial population. Dot size indicates gene count and color indicates adjusted p-value. No significantly enriched KEGG pathways were identified for the homeostatic population.

**Figure S5:**
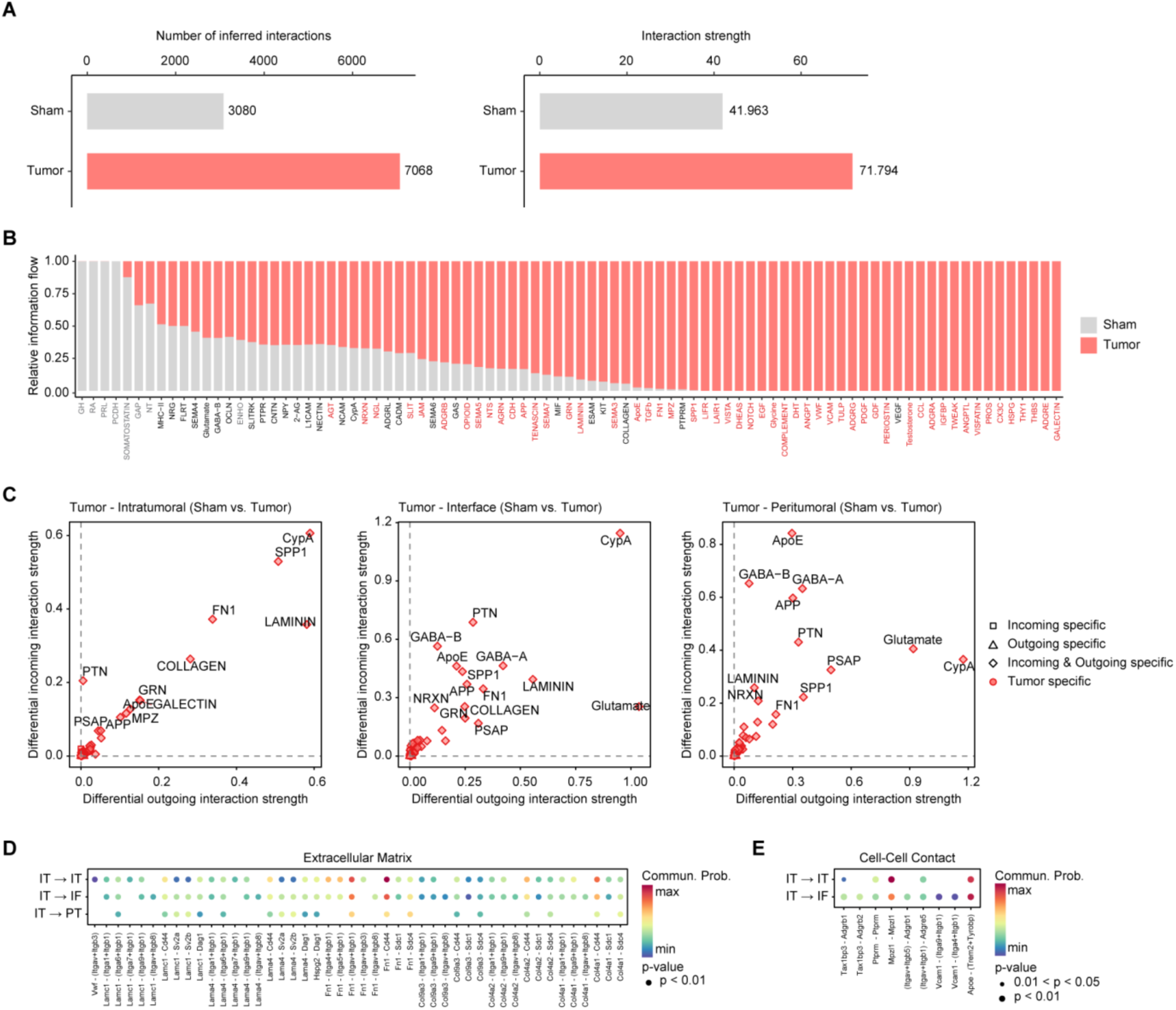
Brain metastasis remodels spatial intercellular communication networks. A) CellChat comparison of the total number and aggregate strength of inferred cell-cell communication interactions in tumor-bearing and sham brains. B) Relative information flow of inferred signaling pathways in tumor-bearing and sham brains. Pathways are ordered according to their relative contribution across the two conditions; pathways highlighted in red are significantly enriched in tumor-bearing samples by Wilcoxon test (p < 0.05). C) Incoming and outgoing signaling strength across intratumoral, interface, and peritumoral spatial domains. Each point represents an inferred signaling pathway; position reflects differential outgoing and incoming interaction strength, and symbol denotes whether the pathway is incoming-specific, outgoing-specific, altered in both directions, or tumor-specific. D-E) Predicted extracellular-matrix ligand-receptor interactions (D) and cell-cell contact interactions (E) originating from the intratumoral domain and directed toward intratumoral (IT→IT), interface (IT→IF), or peritumoral (IT→PT) regions. Dot color represents predicted communication probability, and dot size denotes statistical significance as indicated.

**Figure S6:**
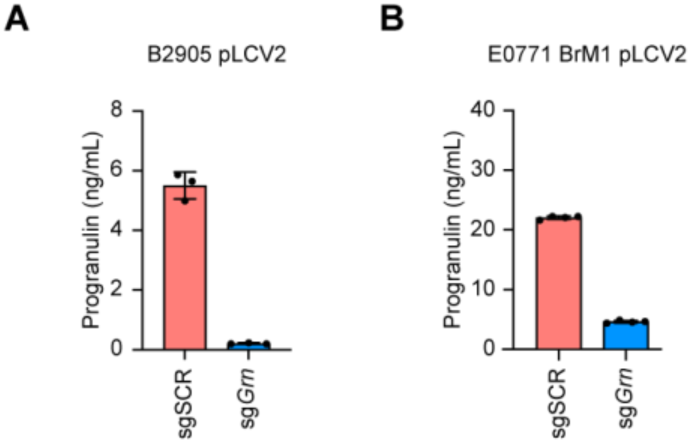
CRISPR-mediated *Grn* knockout reduces PGRN secretion by brain-tropic tumor cells. A-B) PGRN concentrations in conditioned media collected from control sgSCR and *Grn*-targeting sg*Grn* transduced B2905 (A) and E0771 BrM1 (B) cells, measured by ELISA and normalized to cell density. Individual points represent replicate measurements; bars show mean ± SD.

**Figure S7:**
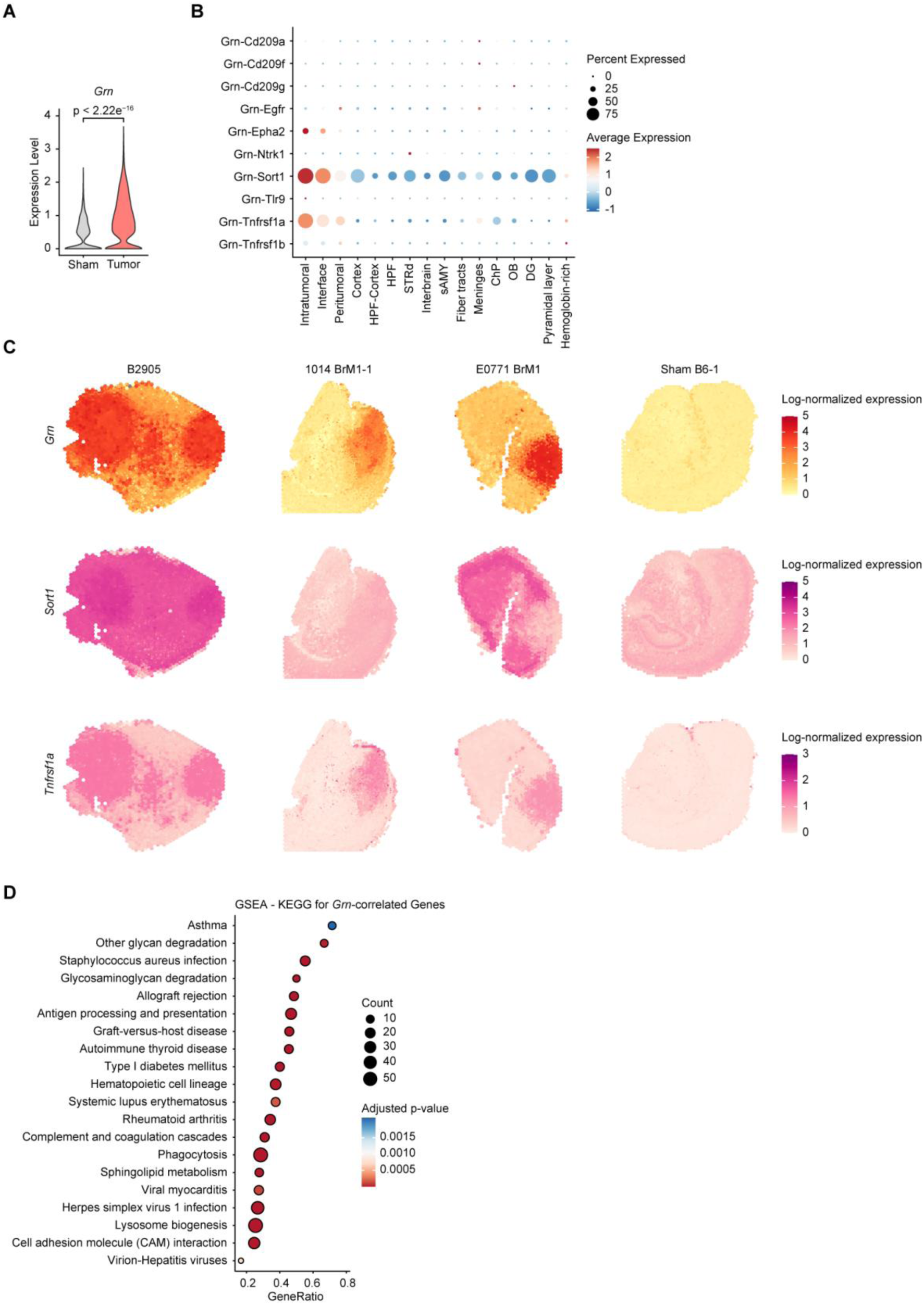
Mouse brain metastases exhibit spatially localized *Grn* expression and candidate PGRN-receptor interactions. A) Comparison of *Grn* expression between tumor-bearing and sham brains in the mouse spatial transcriptomic dataset. P-value calculated using Wilcoxon rank sum test. B) Candidate *Grn*-receptor interactions inferred across spatial tissue domains using NICHES and LIANA. C) Representative spatial feature plots showing log-normalized expression of *Grn*, *Sort1*, and *Tnfrsf1a* in B2905, 1014 BrM1, and E0771 BrM1 tumor-bearing brains and a representative sham brain. D) KEGG gene-set enrichment analysis of genes positively correlated with *Grn* in mouse BM-associated microglia.

**Figure S8:**
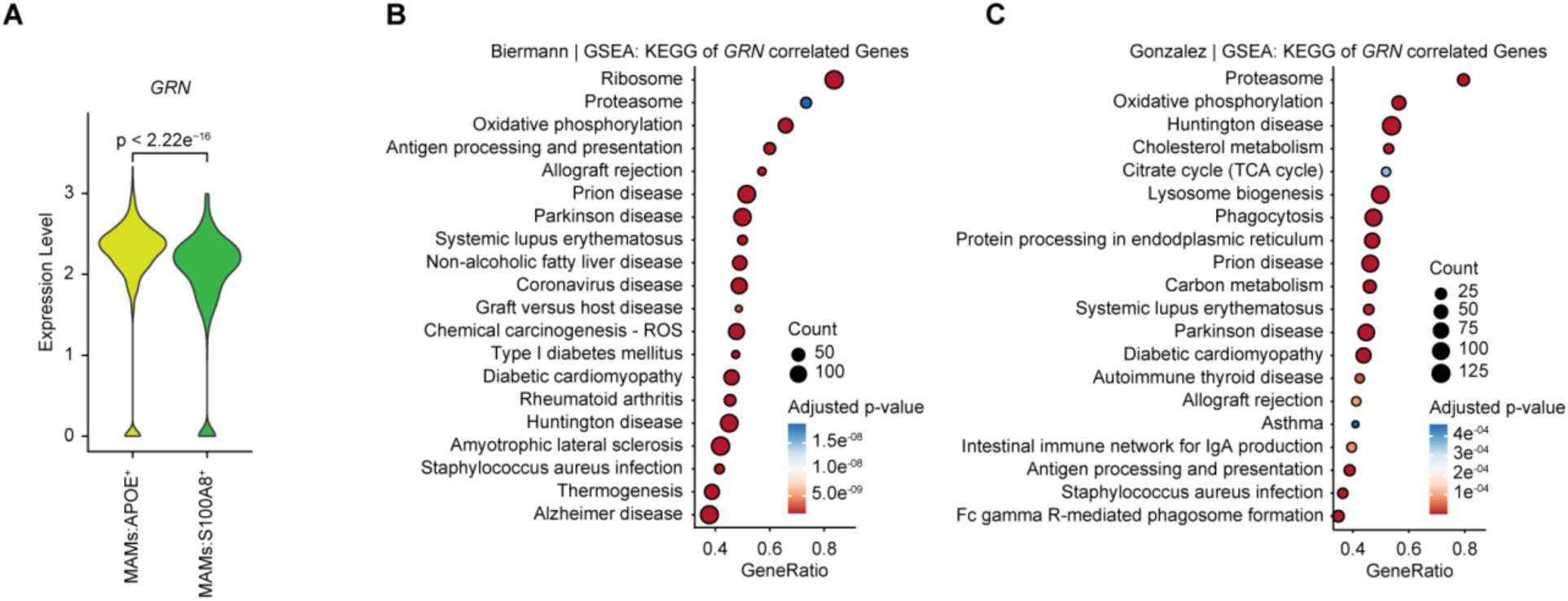
*GRN* is enriched in APOE^+^ metastasis-associated macrophages and correlates with immune and metabolic programs in human brain metastases. A) Comparison of *GRN* expression between APOE^+^ and S100A8^+^ metastasis-associated macrophage (MAM) populations in the Gonzalez et al. human brain metastasis scRNA-seq dataset. P-value calculated using Wilcoxon rank sum test^9^. B-C) KEGG gene-set enrichment analysis of genes positively correlated with *GRN* in combined macrophage and microglia populations from the Biermann et al. melanoma brain metastasis dataset^7^ (B) and the Gonzalez et al. multi-tumor brain metastasis dataset^9^ (C).

**Figure S9:**
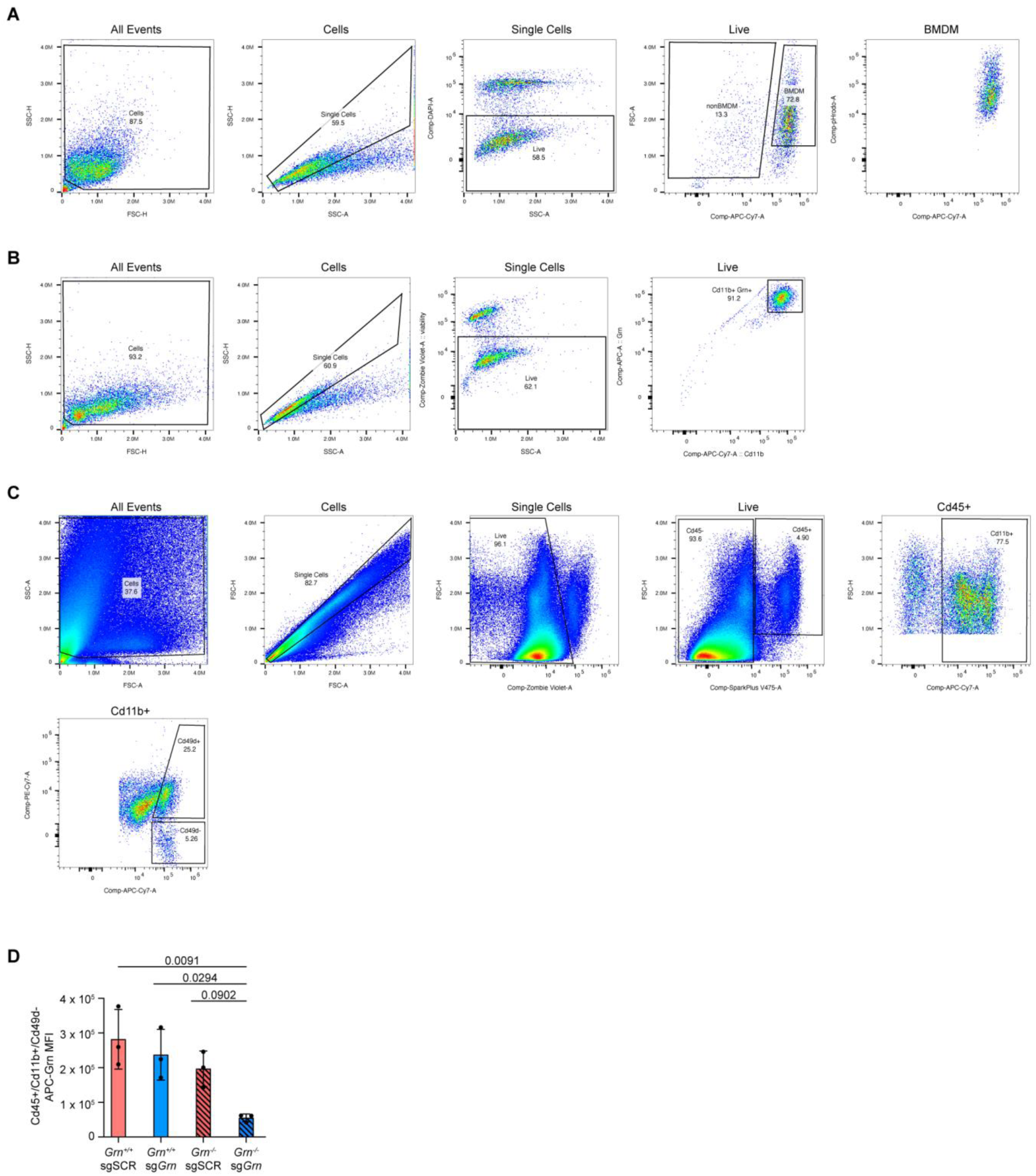
Flow cytometric assessment of tumor-cell engulfment and PGRN abundance in macrophages and microglia. A) Representative flow cytometry gating strategy for quantifying engulfment of pHrodo-labeled tumor cells by bone marrow-derived macrophages (BMDMs). B) Representative flow cytometry gating strategy for quantifying intracellular PGRN in *Grn*^+/+^ and *Grn*^−/−^ BMDMs following coculture with sgSCR or sg*Grn* B2905 tumor cells. C) Intracellular PGRN abundance in microglia isolated from brain metastases generated by intracardiac injection of control sgSCR or *Grn*-deficient sg*Grn* B2905 tumor cells into *Grn*^+/+^ or *Grn^−^*^/−^ hosts. Microglia were gated as live CD45+CD11b+CD49d-cells. D) Quantification of intracellular PGRN by APC mean fluorescence intensity (MFI) in microglia isolated from brain metastases generated by intracardiac injection of control sgSCR or *Grn*-deficient sg*Grn* B2905 tumor cells into *Grn*^+/+^ or *Grn^−^*^/−^ hosts, gating shown in (C). Points represent individual mice, and bars show mean ± SD. Statistical comparisons were performed using one-way ANOVA with Tukey’s multiple comparisons test comparing the mean of each group to every other group.

**Figure S10:**
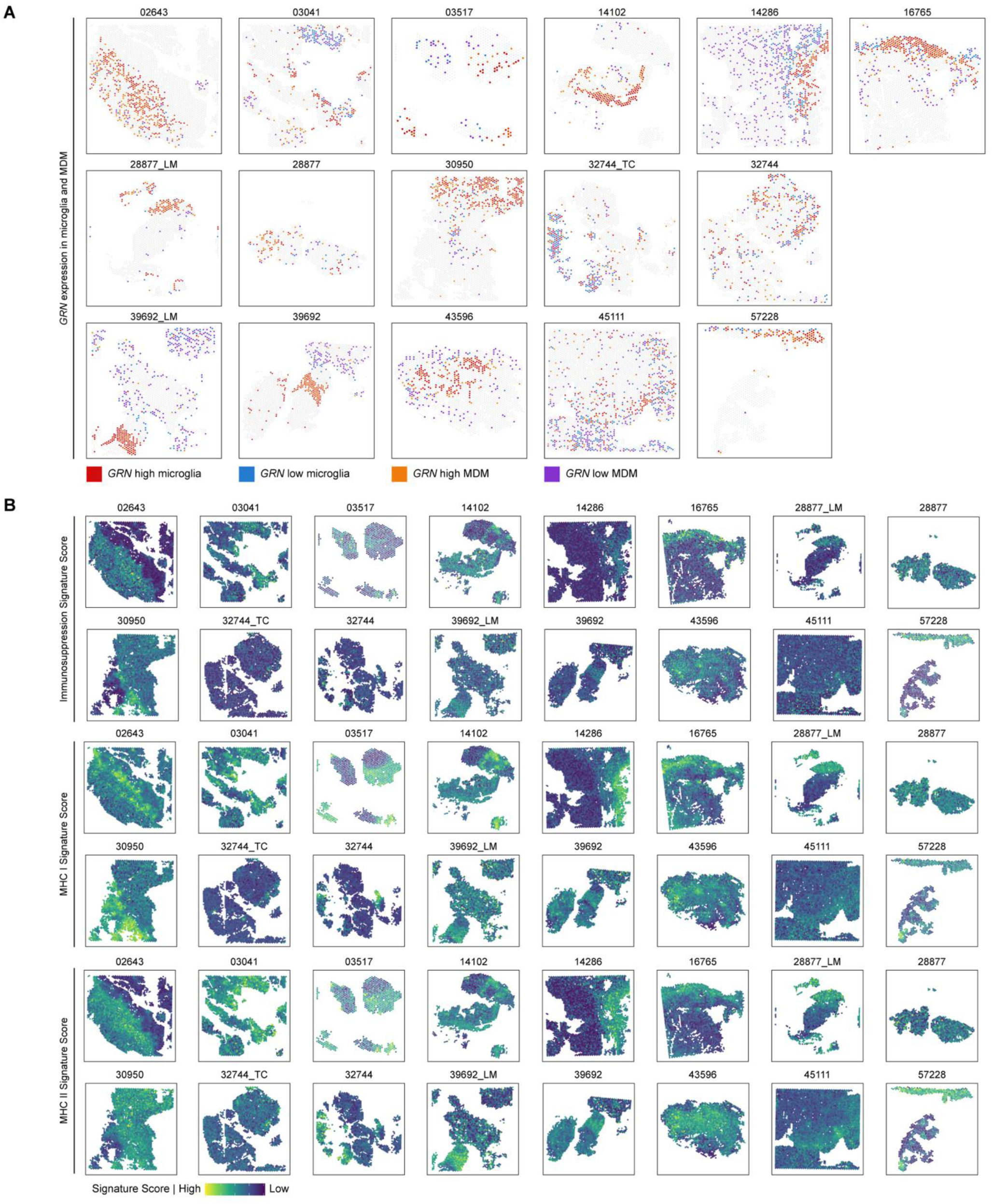
Spatial mapping of *GRN*-high and *GRN*-low myeloid niches and immunoregulatory transcriptional programs in human melanoma brain metastases. A) Spatial distribution of spots enriched for microglial or monocyte-derived macrophage (MDM) transcriptional signatures across 16 human melanoma brain metastasis specimens representing 13 patients. Signature-enriched spots were classified within each specimen as *GRN*-high or *GRN*-low based on the upper and lower 20th percentiles of *GRN* expression, respectively. Red, *GRN*-high microglia; blue, *GRN*-low microglia; Orange, *GRN*-high MDM; Purple, *GRN*-low MDM. Sample identifiers are indicated above each plot. B) Spatial distributions of immunosuppression (top) signature score, and antigen presentation signature scores including MHC class I antigen-presentation (middle), and MHC class II antigen-presentation (bottom) across the same samples. Color scales indicate relative signature scores within each spatial dataset.

**Figure S11:**
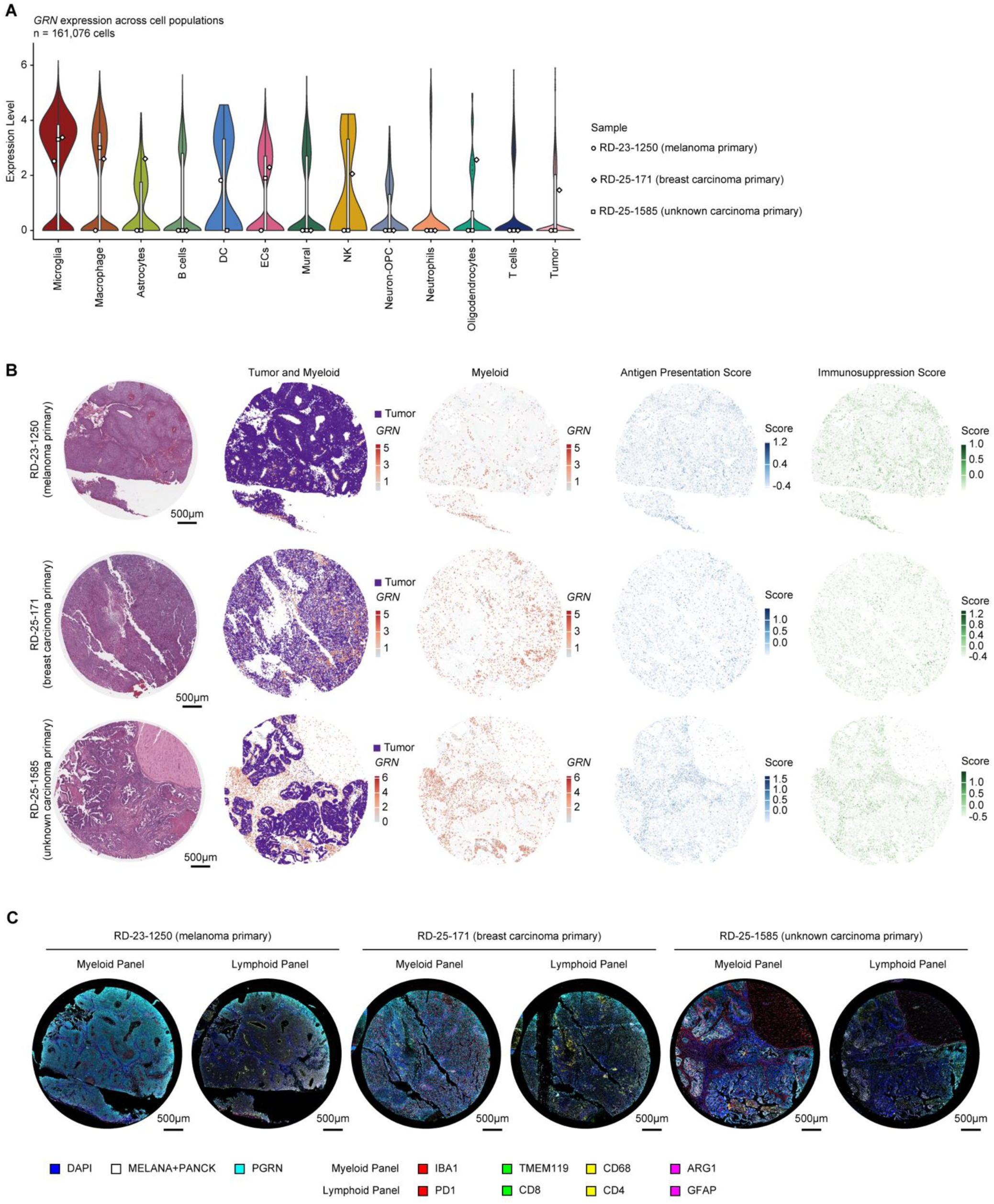
Single-cell spatial profiling associates *GRN*-high myeloid populations with immunoregulatory programs in human brain metastases. A) Violin plots showing *GRN* expression across annotated cell populations in Xenium spatial transcriptomic data from three human brain metastasis specimens. Shapes indicate median expression for each individual specimen. A total of 161,076 cells were analyzed. MDM, monocyte-derived macrophage; DC, dendritic cell; ECs, endothelial cell; OPC, oligodendrocyte precursor cell. B) Representative spatial maps from the three brain metastases showing hematoxylin and eosin (H&E) staining, annotated tumor and myeloid populations, *GRN* expression within myeloid cells, antigen-presentation signature scores, and immunosuppression signature scores. Specimen identifiers are shown at left. C) Multiplex immunofluorescence staining of the same three human brain metastases analyzed using Xenium spatial transcriptomics.

**Table S1:** Immune cell gene signatures derived from mouse BM scRNA-seq dataset.

| **Microglia** | **MDM** | **BAM** | **Neutrophils** | **DC** | **NK** | **T cells** |
| --- | --- | --- | --- | --- | --- | --- |
| *Hpgds* | *Sirpb1c* | *Cbr2* | *S100a8* | *Kmo* | *Klre1* | *Ptprcap* |
| *Ltc4s* | *Apoc2* | *Cd163* | *Cxcr2* | *Ffar4* | *Klrb1c* | *Cd3g* |
| *Fcgr1* | *Klra2* | *Lyve1* | *Mmp9* | *Clec9a* | *Nkg7* | *Cd3e* |
| *Tmem119* | *Clec4a1* | *Pf4* | *Retnlg* | *Btla* | *Gzma* | *Cd2* |
| *Cd68* | *Hp* | *F13a1* | *Hdc* | *Gcsam* | *Il2rb* | *Gimap3* |
| *Ccr5* | *F10* | *Clec4n* | *S100a9* | *Xcr1* | *Ncr1* | *Cd3d* |
| *Cd84* | *Chil3* | *Fcna* | *Il1f9* | *Cd209a* | *Txk* | *Lat* |
| *Rab3il1* | *Sirpb1b* | *Cd209g* | *Stfa2l1* | *Itgae* | *Klrc2* | *Trac* |
| *Gpr34* | *Ccr2* | *Mrc1* | *Wfdc21* |  | *Gzmb* | *Lck* |
| *Cx3cr1* | *Cyp4f18* | *Fpr1* | *Ptgs2os2* |  | *Klrb1f* | *Skap1* |
| *Fcgr3* | *Ms4a8a* | *Pla2g2d* |  |  |  | *Itk* |
| *P2ry12* | *Clec4e* | *Ms4a7* |  |  |  | *Cd247* |
| *Crybb1* | *Adgre4* | *Cd209f* |  |  |  | *Gimap7* |
| *Aif1* | *Lilrb4a* | *Clec10a* |  |  |  | *Cxcr6* |
| *Fcrls* |  | *Ms4a14* |  |  |  | *Cd28* |
| *Csf1r* |  |  |  |  |  |  |
| *C1qc* |  |  |  |  |  |  |
| *C1qa* |  |  |  |  |  |  |
| *C1qb* |  |  |  |  |  |  |

**Table S2:**
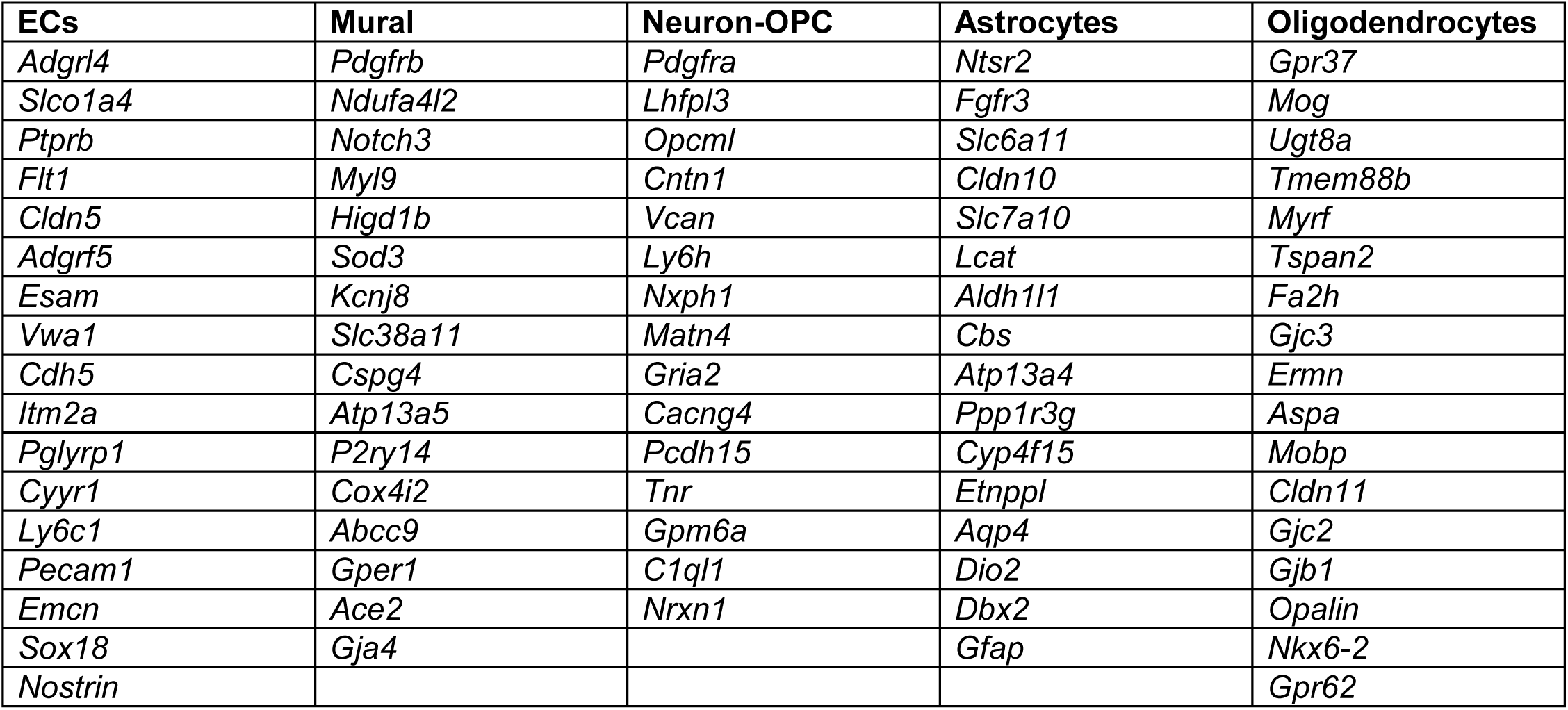
Non-immune cell gene signatures derived from mouse BM scRNA-seq dataset.

**Table S3:**
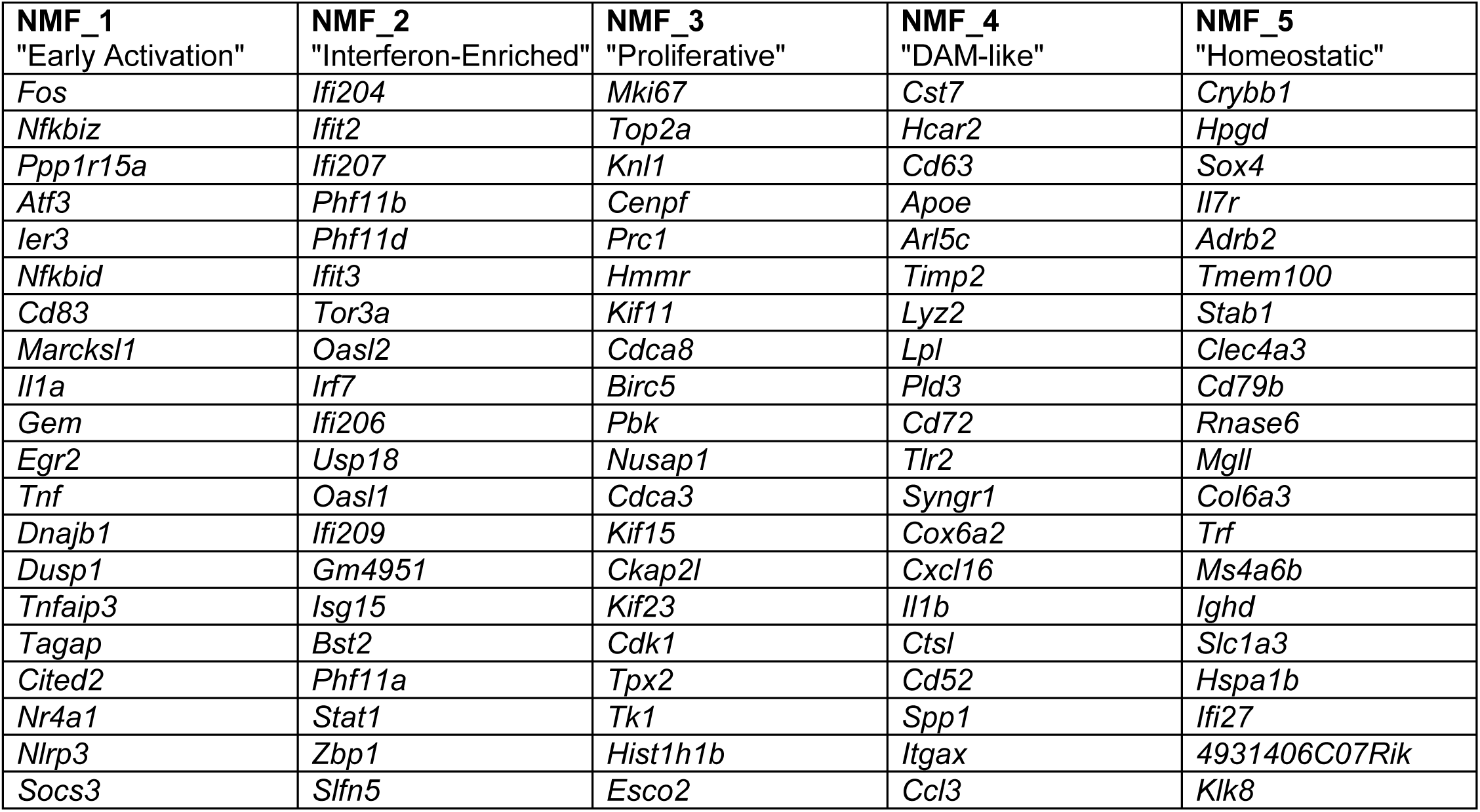
Top 20 genes defining NMF programs.

**Table S4:**
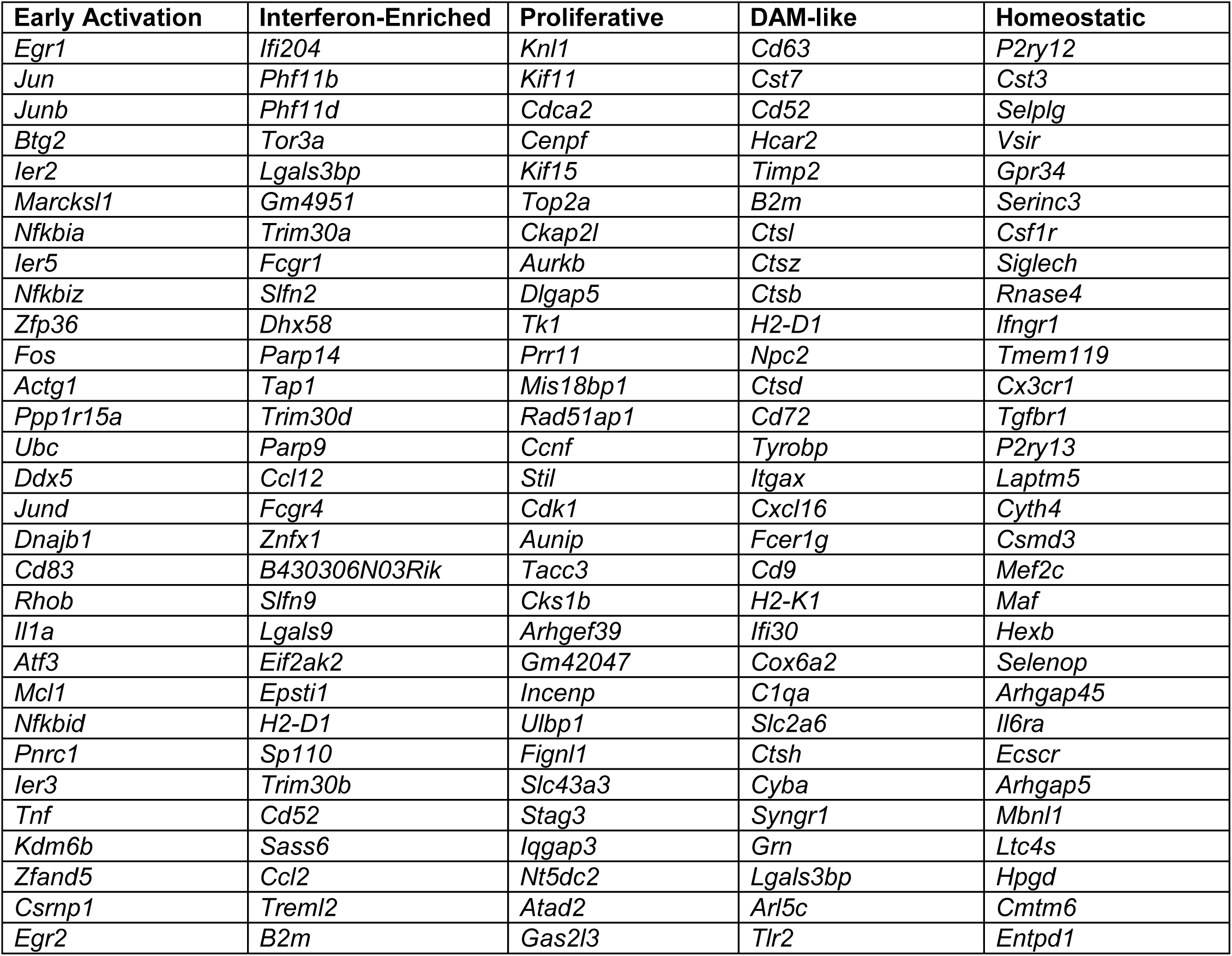
Top 30 marker genes of BM-associated NMF-defined microglial subtypes.

**Table S5:** Human orthologs of immune cell gene signatures derived from mouse BM scRNA seq dataset used in human Visium dataset analysis.

| Microglia | MDM |
| --- | --- |
| *HPGDS* | *SIRBP1* |
| *LTC4S* | *APOC2* |
| *FCGR1* | *CLEC4A* |
| *TMEM119* | *HP* |
| *CD68* | *F10* |
| *CCR5* | *CHIA* |
| *CD84* | *CCR2* |
| *RAB3IL1* | *CYP4F3* |
| *GPR34* | *MS4A8* |
| *CX3CR1* | *CLEC4E* |
| *FCGR2A* | *LILRB4* |
| *P2RY12* |  |
| *CRYBB1* |  |
| *AIF1* |  |
| *FCRL2* |  |
| *CSF1R* |  |
| *C1QC* |  |
| *C1QA* |  |
| *C1QB* |  |

**Table S6:**
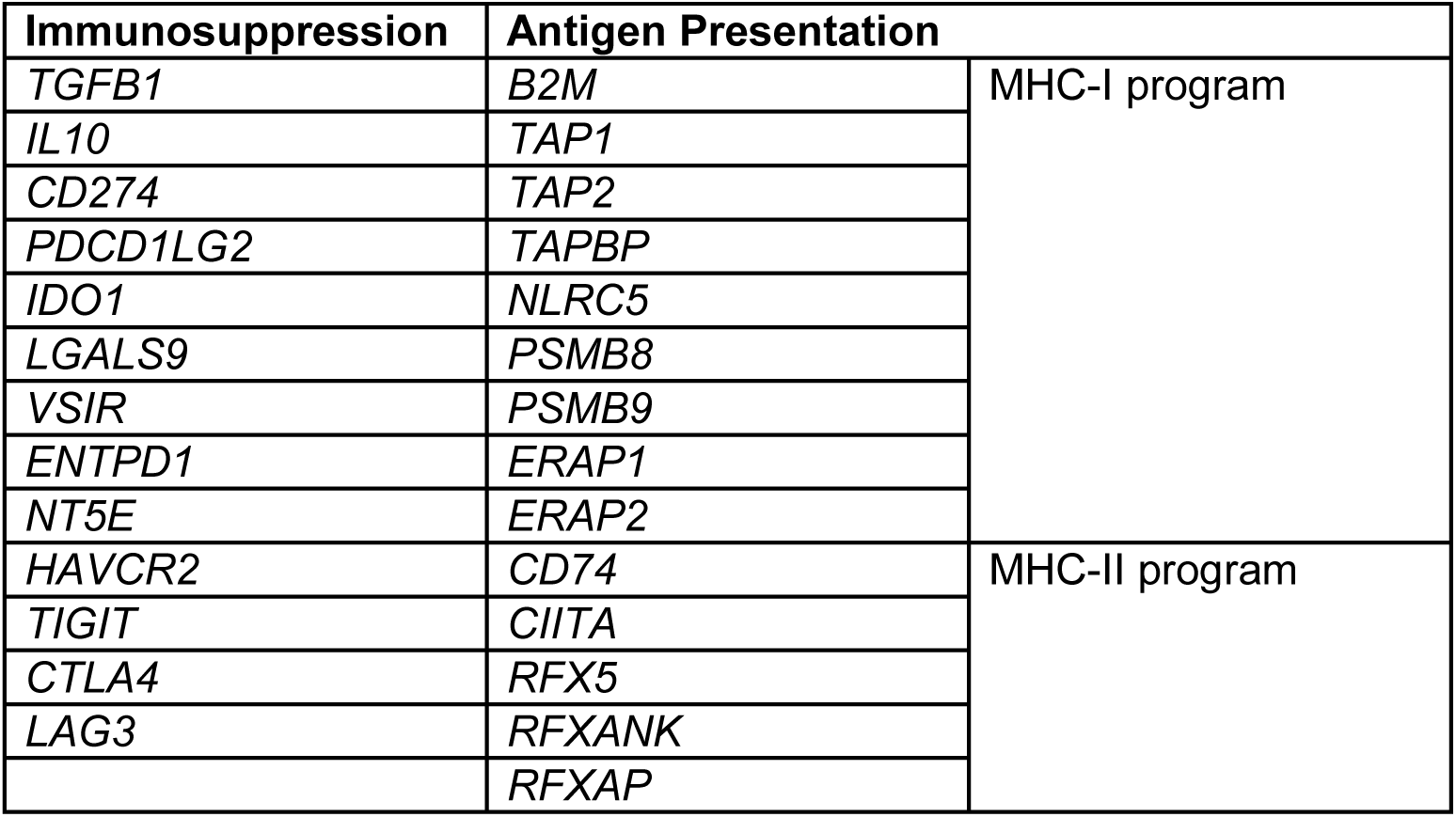
Human immunosuppression and antigen presentation signatures used in human Visium dataset analysis.

| Immunosuppression | Antigen Presentation |  |
| --- | --- | --- |
| *TGFB1* | *B2M* | MHC-I program |
| *IL10* | *TAP1* |  |
| *CD274* | *TAP2* |  |
| *PDCD1LG2* | *TAPBP* |  |
| *IDO1* | *NLRC5* |  |
| *LGALS9* | *PSMB8* |  |
| *VSIR* | *PSMB9* |  |
| *ENTPD1* | *ERAP1* |  |
| *NT5E* | *ERAP2* |  |
| *HAVCR2* | *CD74* | MHC-II program |
| *TIGIT* | *CIITA* |  |
| *CTLA4* | *RFX5* |  |
| *LAG3* | *RFXANK* |  |
|  | *RFXAP* |  |

**Table S7:** Human orthologs of immune cell gene signatures derived from mouse BM scRNA seq dataset used in human Xenium dataset analysis.

| Pan-myeloid | Microglia | MDM |
| --- | --- | --- |
| CD68 | P2RY12 | MRC1 |
| CD14 | CXCR1 | CD163 |
| CSF1R | TMEM119 | VSIG4 |
| FCGR2A | GPR34 | STAB1 |
| CYBB | TREM2 | F13A1 |
| NCF1 | MERTK | MSR1 |
| NCF2 |  | CSF1R |
| LIPA |  | MAFB |
| CIITA |  | FOLRCIITA2 |
|  |  | CCR2 |
|  |  | CD14 |
|  |  | FCGR2A |
|  |  | FCN1 |

**Table S8:** Human immunosuppression and antigen presentation signatures used in human Xenium dataset analysis.

| **Immunosuppression** | **Antigen Presentation** |
| --- | --- |
| *CD163* | *CD80* |
| *MRC1* | *CD86* |
| *ARG1* | *CD83* |
| *ARG2* | *CD40* |
| *IL10* | *CD74* |
| *TGFB1* | *CIITA* |
| *TGFB2* | *NLRC5* |
| *TGFB3* | *TAP1* |
| *HMOX1* | *TAP2* |
| *VEGFA* | *PDIA3* |
| *IL4R* | *CANX* |
| *CCL22* | *ITGAX* |
| *CCL17* | *ITGAM* |
| *CCL18* | *CSF1R* |
| *PPARG* | *FCGR1A* |
| *SOCS3* | *FCGR2A* |
| *STAB1* | *FCGR3A* |
| *FOLR2* | *CCR7* |
| *F13A1* | *CCL19* |
| *LGMN* | *LAMP3* |
| *THBS1* | *XCR1* |
| *MMP9* | *CLEC9A* |
| *MMP12* | *FCER1A* |
| *IL13RA1* |  |
| *IL4* |  |
| *CLEC10A* |  |
| *SLC40A1* |  |
| *CXCL10* |  |
| *PDCD1LG2* |  |
| *CD274* |  |

**Table S9:** Clinical metatadata for human brain metastasis Visium cohort.

| *Sample_ID* | *PatientID* | *ROI Type* | *Gender* | *Age at BrMet Resection* | *Race* | *Ethnicity* | *Focal Radiation* | *WBRT* | *Status* |
| --- | --- | --- | --- | --- | --- | --- | --- | --- | --- |
| 2643 | Patient1 | BTI | Male | 55 | White | Non-Hispanic | Yes | No | Deceased |
| 3041 | Patient2 | LT | Male | 57 | White | Non-Hispanic | Yes | No | Deceased |
| 3517 | Patient3 | BTI | Female | 61 | White | Non-Hispanic | No | Yes |  |
| 14102 | Patient4 | BTI | Female | 68 | White | Non-Hispanic | Yes | No |  |
| 14286 | Patient5 | BTI | Male | 62 | White | Non-Hispanic | No | No | Deceased |
| 16765 | Patient6 | BTI | Female | 75 | White | Non-Hispanic | No | No |  |
| 28877_LM | Patient7 | LT | Male | 18 | Asian | Non-Hispanic | No | No | Deceased |
| 28877 |  | TC |  |  |  |  |  |  |  |
| 30950 | Patient8 | BTI | Male | 75 | White | Non-Hispanic | Yes | No |  |
| 32744 | Patient9 | BTI | Male | 69 | White | Non-Hispanic | Yes | No | Deceased |
| 32744_TC |  | TC |  |  |  |  |  |  |  |
| 39692 | Patient10 | BTI | Male | 49 | White | Non-Hispanic | No | No | Deceased |
| 39692_LM |  | LT |  |  |  |  |  |  |  |
| 43596 | Patient11 | BTI | Female | 51 | White | Hispanic/Latino | No | No | Deceased |
| 45111 | Patient12 | BTI | Male | 65 | White | Non-Hispanic | No | No | Deceased |
| 57228 | Patient13 | TC | Female | 59 | White | Non-Hispanic | Yes | Yes |  |

BTI, Brain-Tumor Interface; LT, Leptomeningeal Tumor; TC, Tumor Center; WBRT, whole brain radiation therapy

## Methods

### Experimental models and methods

#### Cell Culture

B2905, 4T1, and E0771 cells were obtained from ATCC and cultured in RPMI-1640 supplemented with 10% FBS, 1% penicillin-streptomycin and 1% GlutaMAX. HEK293T cells (ATCC) were cultured in Dulbecco’s Modified Eagle Medium (DMEM) supplemented with 10% FBS, 1% penicillin-streptomycin. Yumm1.7 cells (ATCC) were maintained in Ham-F12/DMEM media supplemented with 10% fetal bovine serum (FBS), 1% penicillin-streptomycin, and 1% non-essential amino acids. 1014 cells were generously gifted by Lionel Larue (Institut Curie) and maintained in Ham-F12 media supplemented with 10% FBS, 1% penicillin-streptomycin. Brain-tropic lines generated in this report (Yumm1.7 BrM3, 1014 BrM1, and E0771 BrM1) were maintained in media identical to their respective parental cell lines. All cell lines were maintained in a 5% CO2 incubator at 37°C and were routinely tested for mycoplasma contamination.

#### Mouse models

All experiments were conducted following protocols approved by the NYU Institutional Animal Care Use Committee (IACUC). For studies requiring only wild-type mice, C57BL/6J mice were purchased from Jackson Labs (#000664). For studies requiring *Grn^−/−^*mice, PGRN KO mice were purchased from Jackson Labs (#013175) and bred to wild-type C57BL/6J mice; wild-type and knockout littermates were used in all studies conducting direct comparisons between the two genotypes. Mouse genotyping was performed by Transnetyx. For experiments using 4T1 cells, BALB/cJ mice were purchased from Jackson Labs (#000651). All mice were injected for experiments at 6-12 weeks of age, and both male and female mice were used, ensuring age- and sex- matching across experimental conditions where possible. For all experiments comparing brain metastatic burden across experimental conditions, all mice were euthanized upon weight loss >20% and/or signs of distress (neurological signs, hunching) in any group.

#### Plasmid generation

For CRISPR/Cas9-mediated knockout: sgRNA sequences were designed using the GPP sgRNA Designer (Broad Institute) and cloned into pLentiCRISPRv2-Puro using *BsmBI* digestion followed by sgRNA sequence ligation^85^. Whole-plasmid sequencing was performed to validate cloning prior to downstream applications (Azenta). For GFP-luciferase overexpression: CMV-Luciferase-EF1α-copGFP Lentivector Plasmid was purchased from BD Biosciences (BLIV511PA-1).

#### Lentiviral Production and Infection

HEK293T cells at 80% confluency were co-transfected with 12 μg of lentiviral expression constructs, 8 μg of psPAX2 and 4 μg pMD2.G vectors using Lipofectamine 2000 (LifeTech 11668019) following manufacturer’s recommendations. At 48 and 72 hr post transfection, viral supernatants were collected, filtered (0.45 μm) and stored at −80°C. Cells were infected with lentiviral supernatant supplemented with polybrene (4 ug/mL) followed by antibiotic selection 48 hours after infection. For selection with pLentiCRISPRv2-Puro plasmids, cells were maintained as pools in 2 ug/mL puromycin. For selection with GFP-luc plasmids, cells were sorted on a SY3200 Cell Sorter.

#### In vitro proliferation assay

Proliferation assays were performed using the Incucyte Live-Cell Analysis System (Sartorius). A total of 5,000 cells were seeded per well in 96-well plates and incubated overnight to allow for adherence. Plates were then transferred to the Incucyte, where cells were monitored for at least 72 hours. Brightfield whole-well images were acquired every 8 hours using Incucyte Software V2023.1, and percent confluency per well was quantified over time. For experiments involving recombinant mouse progranulin (rmPGRN), rmPGRN or vehicle control was added following the overnight adhesion period, immediately prior to initiation of Incucyte imaging.

#### Bone marrow-derived macrophage culture & RNA sequencing

Bone marrow was isolated from the femurs of 8-10 week old male *Grn*^+/+^ and *Grn*^−/−^ littermates. Marrow cells were recovered by centrifugation, pelleted at 1,200 rpm for 5 min at 4°C, and treated with ACK lysis buffer for 1 min to remove erythrocytes. Following quenching with PBS, cells were passed through a 70-µm filter, pelleted, and cultured in DMEM GlutaMAX supplemented with 20% heat-inactivated fetal bovine serum, 1% penicillin-streptomycin, and 100 ng/mL MCSF (R&D Systems). Approximately 4 × 10⁶ cells were plated per well in six-well plates and maintained at 37°C with 5% CO₂. Nonadherent cells were removed and the differentiation medium was refreshed one, three and five days after isolation. On day 6, differentiated bone marrow-derived macrophages (BMDMs) were detached using 500 µL CTS Versene per well for 5 min at 37°C, gently scraped into suspension, and replated at 5 × 10⁵ cells per well in 12-well plates. Three replicate wells were prepared for each genotype. After 6 hours, the medium was replaced with 0.5 mL serum-free DMEM containing antibiotics, 100 ng/mL MCSF, and 100 ng/mL LPS. The following day, cells were placed on ice, washed with PBS, and lysed directly in each well using 200 µL DNA/RNA Shield (Zymo Research). Lysates were submitted to Plasmidsaurus for RNA extraction, library preparation, and 3′-end-counting RNA sequencing on an Illumina NovaSeq X Plus. Reads were processed using the Plasmidsaurus RNA-seq analysis pipeline and aligned to the *Mus musculus* GRCm39 reference genome, release 114. Gene-level quantification and differential-expression analysis comparing *Grn*^+/+^ and *Grn*^−/−^ BMDMs were performed using the Plasmidsaurus analysis platform.

#### pHrodo tumor-cell uptake assay

*Grn*^+/+^ and *Grn*^−/−^ BMDMs were generated as described above and cocultured for 24 h with control sgSCR or *Grn*-deficient sg*Grn* B2905 tumor cells labeled using the Incucyte pHrodo Red Cell Labeling Kit (Sartorius, catalog no. 4649). Following coculture, cells were washed once with PBS, gently scraped into suspension, transferred to V-bottom plates, and pelleted at 1,200 rpm for 4 min at 4°C. Cells were incubated in 50 µL Fc-blocking solution containing 0.5 µg/mL anti-mouse CD16/32 antibody for 10 min on ice and subsequently stained with APC/Cy7-conjugated anti-CD11b antibody and DAPI for 20 min on ice. Cells were washed twice with flow-cytometry buffer, resuspended in 150 µL buffer, and analyzed using a Cytek Aurora. Live BMDMs were identified as DAPI-CD11b+ cells, and tumor-derived pHrodo signal was quantified as mean fluorescence intensity within this population using FlowJo v10.8.1 software.

#### Progranulin ELISA

Progranulin concentrations in conditioned media were measured using a mouse progranulin ELISA kit (R&D Systems) according to the manufacturer’s instructions. Media was collected from cultured cells after 24-72 hours and centrifuged at 1,000 × *g* for 4 minutes to remove cells and debris. A 50 μL aliquot of clarified media was used per sample for measurement by ELISA. Measured concentrations were normalized to cell density at the time of collection, as determined by percent confluency using the Incucyte imaging system.

#### Intracardiac injections

Prior to injection, mice were anesthetized in an induction chamber with an isoflurane vaporizer. Following anesthesia, the thorax of each animal was shaved with a razor blade and cleansed with 3 applications of 10% povidone-iodine alternating with 3 applications of isopropyl alcohol. An ultrasound (Visualsonics Vevo 770 Ultrasound Imaging System) was used to visualize the left ventricle, and a stereotactic injector was used to inject 100,000 cells suspended in 100 μL PBS with continued ultrasound guidance.

#### Intracarotid injections

Intracarotid injections were performed as previously described^86^. Briefly, animals were anesthetized by ketamine (100 mg/kg) and xylazine (10 mg/kg). The procedure area was shaved with a razor blade and cleaned with 3 applications of 10% povidone-iodine alternating with 3 applications of isopropyl alcohol. Each animal was positioned under a stereo microscope on a warming pad for the procedure. After incising the skin from half the neck down to the sternum, the common carotid artery was isolated for injection, with ligatures pre-positioned to control post-injection bleeding. For each animal, 100,000 cells suspended in 100 μL of PBS were injected. Mice were monitored daily postoperatively and administered subcutaneous buprenorphine every 12 h for 72 h post-surgery at a concentration of 0.1 mg/kg.

#### Tissue processing for FFPE

Mice exhibiting >20% weight loss and/or neurologic symptoms were euthanized by carbon dioxide inhalation at 4L/min for 4 minutes, followed by cervical dislocation. Tissues were dissected and processed as previously described^87^. Organs bearing metastases (brains, livers, kidneys and adrenal glands) were fixed in 10% neutral buffered formalin for 72 hours. Organs were then washed three times in PBS and dehydrated in 70% ethanol for 24 hours. Brains were grossly divided into coronal thirds, and livers were sectioned by lobe. All tissues were paraffin-embedded and 5 μm sections were generated. For brains, each embedded coronal third was serially sectioned at 50 μm intervals. Two evenly spaced levels were selected from each third, yielding a total of six coronal brain section levels per sample for metastatic quantification. For livers, 5 μm sections were collected from two levels spaced 100 μm apart. All sections were stained with hematoxylin and eosin (H&E).

#### Quantification of metastatic burden

H&E-stained sections were scanned at a 40X magnification (pixel size 0.25 µm) on a Leica AT2 whole slide scanner and the image files uploaded to the OMERO Plus image data management system (GlencoeSoftware). Metastasis quantification was then performed using HALO, a digital pathology image analysis platform that uses AI for annotation of tissue classifiers (Indica Labs). Brain metastasis burden for each mouse was quantified as the percent of tumor-occupying area across the six visualized brain tissue sections.

#### Multiplex immunofluorescence

Multiplex immunofluorescence was performed using Akoya Biosciences Opal reagents. Sections were immunostained on a Leica BondRx auto-stainer according to the manufacturer’s instructions. In brief, sections were deparaffinized, followed by antigen retrieval with ER2 (Leica, AR9640; pH9) retrieval buffer at 100° for 20 minutes. Sections were then treated with 3% H_2_O_2_ to inhibit endogenous peroxidases. After blocking with Primary Antibody Diluent (Leica, AR93520), slides were incubated with the first primary antibody and secondary HRP polymer pair, followed by HRP-mediated tyramide signal amplification with a specific Opal® fluorophore. Once the Opal® fluorophore was covalently linked to the antigen, primary and secondary antibodies were removed with a heat retrieval step. This sequence was repeated 5 more times with subsequent primary and secondary antibody pairs, using a different Opal fluorophore with each primary antibody (see Key Resources table for primary antibody and reagent details). After antibody staining, sections were counterstained with spectral DAPI (Akoya Biosciences, FP1490) and mounted with ProLong Gold Antifade (ThermoFisher Scientific, P36935). Semi-automated image acquisition was performed on a PhenoImager HT (previously Vectra® Polaris) multispectral imaging system from Akoya Biosciences. An initial scan at 20x magnification was performed to establish the appropriate exposure settings. Then, using annotations from the initial scan, a spectral unmixing library was generated with HT software (v 2.0.0) and InForm® (V 3.0). This library was used to create an “unmixed” scanning protocol. Slides were then imaged a second time at 20x magnification with the unmixed protocol, producing spectrally unmixed whole slide qptiff scans. Whole-slide scans were imported into HALO AI, version 4.0 (Indica Labs). Nuclei were segmented using DAPI, and cell populations were phenotyped and quantified with the Multiplex IHC module using predefined marker-intensity thresholds. Tumor regions were delineated based on model-specific tumor-marker expression (*Melana* for melanoma models and *Krt8* for breast cancer models) together with DAPI-defined nuclear density and morphology. Within each annotated tumor region, HALO calculated the percentage of cells positive for each marker, and these values were exported directly for downstream analysis.

#### Flow cytometric assessment of mouse brain immune cells

Brains grossly dissected from euthanized mice were briefly washed in HBSS prior to being transferred to a 10-cm plate. Cerebrums were separated from the cerebellums, which were discarded. Cerebrums were manually minced using a razor blade followed by incubation with DNAse and collagenase for 45 minutes at 37 °C with constant shaking. Dissociated tissues were filtered through a 70μM, then a 40μM filter and spun at 300 x g for 10 min. The resulting pellet was resuspended in 38% Percoll and spun at 800 x g for 15 minutes to remove myelin debris. After debris removal, cells were resuspended in 500 μL ACK lysis buffer for 1 minute, then washed with PBS. Cells were resuspended in PBS and counted to aliquot 2M cells per sample. Each sample was resuspended in Zombie Violet cell viability stain diluted in PBS for 10 minutes on ice. Following the viability stain, cells were washed with FACS buffer and spun at 300 x g for 5 min followed by resuspension in 50uL FACS buffer containing 1ug Fc-blocking solution containing anti-mouse CD16/32 antibody and incubated for 15 min on ice. Cells were then directly stained with 50 uL FACS buffer containing fluorophore-conjugated antibodies for 30 min on ice. Cells were washed in FACS buffer before analysis on a Cytek Aurora. Data was analyzed using FlowJo v10.8.1 software.

#### Intracellular PGRN quantification

Cells from whole mouse brains or BMDM co-cultures were prepared as described above. For experiments assessing intracellular PGRN, following cell surface staining, cells were fixed with 4% PFA for 10 min at room temperature, permeabilized with 2% BSA + 0.5% Triton X100, and stained with APC-conjugated anti-PGRN antibody (R&D Systems) for 30 minutes at room temperature. Samples were read on a Cytek Aurora. Data was analyzed using FlowJo v10.8.1 software.

#### Statistical Analysis

Statistical analyses for in vitro and in vivo experiments were performed with Prism 10 (GraphPad Software), with specific tests indicated in figure legends. Unless otherwise stated, displayed bar graph values are averages ± standard deviation.

### Computational Data Generation, Processing, and Analysis

#### Public scRNA-seq datasets of mouse and human brain metastasis

Human scRNA-seq datasets were retrieved from the Gene Expression Omnibus (GSE200218, GSE186344)^7,9^. Cell type annotations were derived from accompanying metadata provided by the original studies. Gene expression among cell types was analyzed and visualized with Seurat. Mouse scRNA-seq from Rodriguez-Baena et al. was obtained from the Gene Expression Omnibus (GSE263623).106 We performed our independent analysis in Seurat. We excluded cells with fewer than 1,000 or more than 5,000 detected genes, or with >10% mitochondrial transcript content, log-normalized the dataset, and generated clusters from the top 30 principal components with a resolution of 0.5. We then generated a subset with only the microglial cells to identify microglial programs and subtypes via NMF (using rank = 5).117 Gene signatures of the major cell types and microglial subtypes were curated using FindAllMarkers(), and these signatures were mapped to our spatial transcriptomics dataset with the AddModuleScore() function. Visualization of these signatures was performed in subspot resolution with BayesSpace as described above. Monocle3 was used to measure pseudotime on the microglial subset as previously described^88^.

### For mouse spatial transcriptomic studies

#### Mouse tissue collection and Visium processing

Tumor-bearing and sham control brains were processed as described above; however, in lieu of formalin fixation and paraffin embedding, brain tissues were snap-frozen in optimal temperature cutting compound (OCT). Cryosections were cut at 10 μm thickness and mounted onto Visium Spatial Gene Expression slides. Tissue sections were warmed, dehydrated, and stained with H&E, followed by brightfield imaging to assess and annotate histological features. The Visium Spatial Gene Expression Reagent Kit (10x Genomics) was then used for permeabilization, reverse transcription, second-strand synthesis, and cDNA amplification. Libraries were sequenced on an Illumina Novaseq X+, and reads were mapped to the mm10 mouse reference transcriptome using Space Ranger v1.3.1 (10x Genomics).

#### Preprocessing, quality control, and tissue domain identification

The filtered_feature_bc_matrix.h5 output files generated by Space Ranger were loaded into Seurat for initial per-sample processing^89^. Spatial Seurat objects were filtered to retain spots with at least 1000 UMIs (unique molecular identifiers). Gene expression matrices were log-normalized, and principal component analysis (PCA) was performed on the top 5000 variable genes. The top 30 principal components were selected for downstream clustering of the individual samples. For multi-sample analyses, Seurat objects for individual samples were merged into a single Seurat object. The merged object was input into BANKSY (using lambda 0.1 and k_geom = 6) and then Harmony for spatially aware clustering and batch correction, respectively^20,21^. Clusters were visualized with Seurat’s SpatialDimPlot() function and manually annotated by tissue region, based on spatial context and marker gene expression. Final marker genes of tissue domains were generated using the FindAllMarkers() function.

#### Cell type deconvolution and probing

To estimate the cellular composition of each spatial spot, we used STdeconvolve, an unsupervised method based on latent Dirichlet allocation (LDA)^23^. The number of topics (cell types) was set to K = 20, selected based on model perplexity and biological interpretability. Marker genes (based on log2 fold change) for each deconvolved topic were compared to reference single-cell RNA-seq datasets (via https://linnarssonlab.org/adolmouse/ and https://brainimmuneatlas.org/) to assign putative cell type identities.

#### Ligand-receptor interaction analysis

To assess spatially localized intercellular communication, we used CellChat, NICHES, and LIANA^44,65,66^. CellChat was applied to identify overrepresented ligand-receptor interactions within spatial clusters. Two CellChat objects, one comprised of all tumor samples and one with all sham samples, were merged for comparative analyses. Signaling pathways were inferred using built-in reference databases. LIANA was used to compile candidate *Grn*-receptor interactions across multiple curated resources. These interactions were supplied to NICHES as a custom ligand-receptor database, and the NeighborhoodToCell output from RunNICHES() was used to estimate spatial interaction scores based on *Grn* expression in neighboring Visium spots and receptor expression in receiving spots across the mouse BM atlas.

#### Subspot resolution feature mapping

We applied BayesSpace to increase spatial resolution by estimating subspot-level gene expression patterns^90^. BayesSpace::spatialEnhance() was run with q = 30 clusters and d = 30 principal components to generate enhanced-resolution feature maps for select genes and deconvolved cell types. Enhanced plots were visualized using featurePlot(), using inferred logcounts from subspots for individual features, or the sum of the inferred logcounts when visualizing multi-gene sets.

#### GRN correlation and pathway-enrichment analyses

Gene-correlation analyses were performed within the designated spots or myeloid populations from each dataset, as specified in the Results section. Log-normalized expression values were used to calculate Pearson correlation coefficients across individual spots/cells between the expression of *Grn* (mouse) or *GRN* (human) and every other detected gene. Genes with undefined correlation coefficients were excluded, and *Grn/GRN* itself was removed before genes were ranked by decreasing correlation coefficient. The most positively correlated genes in each dataset were visualized by heatmap. Expression values were standardized to row-wise Z-scores, and genes were hierarchically clustered using Euclidean distance and complete linkage.

For pathway enrichment analyses, the complete gene list was ranked by Pearson correlation coefficient; this ranking was then applied for GSEA with either clusterProfiler (KEGG) or fgsea (Reactome, via MSigDB)^91,92^. Enriched pathways were visualized using ggplot2. KEGG enrichment via clusterProfiler was also utilized for analysis of microglial subtypes identified during our independent analysis of the Rodriguez-Baena et al. scRNA-seq dataset; for this analysis, statistically significant microglia subtype markers from the FindAllMarkers() output (adjusted p-value < 0.05) were ranked according to their log2 fold change.

#### Visualization and statistical analysis

Plots were generated using ggplot2, ComplexHeatmap, and Seurat-based visualization functions unless otherwise specified^93,94^. Differential expression analyses, heatmaps, dot plots, and ligand-receptor networks were generated using standard visualization pipelines with adjustments for publication clarity. Statistical comparisons for differential expression analyses were performed using the Wilcoxon rank-sum test, with p-values adjusted using the Benjamini-Hochberg method unless otherwise stated.

### For human spatial transcriptomic Visium studies

#### Human biospecimen collection and cohort construction

All human subject research was approved by the Stanford Institutional Review Board (IRB-34363) and conducted in accordance with the Declaration of Helsinki. Written informed consent was obtained from all patients. The Stanford STRIDE database was queried for patients (>18 years) treated at Stanford Health Care between 2008 and 2023 with radiographically confirmed brain metastases arising from melanoma. Medical records were individually reviewed to confirm diagnosis and brain metastasis status. Formalin-fixed paraffin-embedded (FFPE) surgical specimens were obtained from Stanford Neuropathology archives and reviewed by brain pathologist (D.S.) to confirm diagnosis and tissue quality. Cohort characteristics are summarized in Supplementary Table 9. Clinical data extracted from electronic health records included gender, age, dates of surgical resections, and focal radiation and/or whole brain radiation status (if applicable).

#### Histopathologic review and region-of-interest classification

16 H&E-stained tissue sections from 13 unique human melanoma brain metastasis patients were reviewed microscopically in consultation with a neuropathologist to identify and classify the spatial regions represented in the Visium dataset. Each of the 16 analyzed tissue sections was assigned to one of three region-of-interest (ROI) categories based on histopathologic morphology: tumor center (TC), defined as an intraparenchymal region composed predominantly of melanoma cells; brain-tumor interface (BTI), defined as an intraparenchymal region in which melanoma and adjacent non-neoplastic brain tissue were directly juxtaposed or intermixed; or leptomeningeal tumor (LM), defined by melanoma involving the leptomeningeal compartment. ROI assignments were made by direct microscopic evaluation of the H&E sections rather than by computational clustering. Clinical leptomeningeal disease status was determined independently from clinical records and was treated as a distinct patient-level variable rather than being inferred from ROI type.

#### Spatial transcriptomic data processing

Spatial transcriptomic data from the 16 human melanoma brain metastasis tissue sections were processed using Space Ranger v2.0.1 (10x Genomics) with the Space Ranger count pipeline. Reads were processed against the GRCh38-2020-A human reference using the Visium Human Transcriptome Probe Set v2.0 GRCh38-2020-A. The same probe-set file was used for all samples. Space Ranger processing incorporated the corresponding H&E tissue image and CytAssist image for each section, together with sample-specific manual Loupe alignment files and Visium slide/capture-area information. The Space Ranger filtered_feature_bc_matrix and spatial image outputs were imported into Seurat in R. For each tissue section, gene-by-spot count matrices were read using Read10X, and spatial image/coordinate information was imported using Read10X_Image. Separate Seurat objects were constructed using the Spatial assay and linked to sample-level metadata.

#### Quality control, spot selection, and normalization

Quality-control metrics included total transcript counts per spot (nCount_Spatial), number of detected features (nFeature_Spatial), and mitochondrial transcript percentage. These metrics were assessed both globally and spatially for each tissue section. Mitochondrial percentage was calculated from genes beginning with MT-when available. Spot filtering was performed in two stages. First, tissue-associated spots were manually selected for each section using Loupe-derived barcode lists, and only barcodes included in the corresponding manually selected tissue region were retained. Second, a common quantitative filtering procedure was applied to the retained spots. Spots were required to contain at least 300 detected features and 500 transcript counts. To remove extreme upper-tail outliers, spots exceeding the sample-specific 99.5th percentile of either nCount_Spatial or nFeature_Spatial were excluded. Filtered sections were independently normalized using Seurat’s NormalizeData function with the LogNormalize method and a scale factor of 10,000. Resulting per-sample LogNormalize Spatial-assay expression data were used in the downstream analyses.

#### Myeloid and immune program scoring

To identify spatial regions enriched for microglial or MDM transcriptional programs, mouse gene signatures were first converted to human orthologs using the babelgene R package. Ortholog mapping required a minimum support value of 3, and the top-supported human ortholog was retained. Gene signatures were then restricted to genes represented in the Visium probe set. The same final gene sets were used across all 16 tissue sections. Microglia, MDM, immunosuppression, and antigen presentation signatures were mapped onto each tissue section, using Seurat’s AddModuleScore on the normalized Spatial assay (Table S5; Table S6). Spots within the top 20% of the within-section module-score distribution were classified as Microglia-high, MDM-high, immunosuppression-high, or antigen-presentation-high.

#### *GRN* expression classification

Normalized *GRN* expression was extracted directly from the Spatial assay rather than represented by a module score. *GRN*-high and *GRN*-low states were defined independently within each tissue section. *GRN*-high spots were defined as spots with positive *GRN* expression at or above the 80th percentile of *GRN*-positive spots within that section. *GRN*-low spots were defined as spots at or below the 20th percentile of *GRN* expression across all spots within the corresponding section.

#### Spatial nearest-neighbor analysis

To determine whether *GRN*-high myeloid-enriched regions were preferentially located adjacent to immunosuppression- or antigen-presentation-high regions, a nearest-neighbor analysis was performed using Visium spot coordinates. For each Microglia-high/*GRN*-high, Microglia-high/*GRN*-low, MDM-high/*GRN*-high, or MDM-high/*GRN*-low center spot, the six nearest neighboring spots, excluding the center itself, were identified. To prevent spatially disconnected regions separated by empty tissue from being treated as immediate neighbors, a sample-specific distance constraint was imposed. The typical Visium spot spacing for each tissue section was defined as the median distance from each spot to its nearest neighboring spot. Candidate neighbors were retained only when their distance from the center spot was no greater than 1.75 times this typical nearest-neighbor spacing. Consequently, center spots could contribute fewer than six neighbors when more distant candidate spots exceeded this threshold. For each center spot, the percentage of retained neighboring spots classified as either immunosuppression-high or antigen-presentation-high was calculated. Center-level percentages were first averaged within each tissue section. For patients represented by more than one tissue section, section-level values were subsequently averaged within each patient so that each patient contributed a single biological replicate.

#### Statistical analysis

*GRN*-high and *GRN*-low neighbor-association measurements derived from the same patients were compared using paired Wilcoxon signed-rank tests. Four prespecified comparisons were tested: Microglia versus immunosuppression, MDM versus immunosuppression, Microglia versus antigen presentation, and MDM versus antigen presentation. Multiple-testing correction across these four comparisons was performed using the Benjamini-Hochberg false-discovery-rate procedure. Tests required at least three patients with paired *GRN*-high and *GRN*-low measurements.

### For human spatial transcriptomic Xenium studies

#### Study Design and Patient Cohort

Three formalin-fixed, paraffin-embedded (FFPE) brain tissue specimens were identified and collected in NYUGSoM under IRB study S16-00122. The samples were processed in tissue-microarray format (TMA) as 3 × 3-mm cores. RD-23-1250 was diagnosed as melanoma, RD-25-171 as breast carcinoma and RD-25-1585 is of unknown origin. In total, 3 cores were analyzed. All spatial transcriptomics experiments were performed using the 10x Genomics Xenium In Situ platform with the Xenium Human Multi-Tissue and Cancer Panel (5,089 target genes) and an additional 100 genes custom panel.

#### Data Processing and Quality Control

Raw Xenium output data was loaded in R (v4.4.2) using a custom workflow that reads count matrices and cell boundary polygons via *pandas* through the *reticulate* interface. TMA core boundaries were defined by polygon coordinates, and each cell centroid was assigned to its cognate donor core using point-in-polygon testing (*sp::point.in.polygon*). Per-cell quality control metrics computed included total transcript count (nCount_Xenium), number of detected genes (nFeature_Xenium), nuclear transcript fraction, cell area (µm^2^), and transcript density. Cells with fewer than 50 transcripts were excluded. Retained counts were log-normalized (LogNormalize, scale factor = 100) using Seurat v5.

#### Dimensionality Reduction and Batch Correction

Count matrices were stored on disk using BPCells. Variable features were selected using a layer-aware approach. Representative cell subsets were selected using LeverageScore-based sketch sampling (*SketchData*; 5,000 cells per sample per disease object). Per-disease sketches were merged and PCA computed on 50 components. Harmony batch correction was applied (*IntegrateLayers*, HarmonyIntegration, grouping by sample_id; 30 PCs retained). Integration quality was assessed by comparing unintegrated versus Harmony-corrected UMAPs. Harmony embeddings were projected to all cells using *ProjectData* (Seurat).

#### Cell Type Annotation

Unsupervised clustering was performed on the Harmony-integrated embedding using a shared nearest-neighbor graph (*FindNeighbors*, k = 20, 30 dimensions) followed by Leiden/Louvain community detection (*FindClusters*) at resolutions 0.2, 0.4, 0.6, and 0.8. Resolution 0.4 was used as the primary annotation resolution (23 clusters). Cluster marker genes were identified by Wilcoxon rank-sum test (*FindAllMarkers*, min.pct = 0.25, log-fold-change threshold = 0.25, positive markers only). Cell types were assigned by integrating: (1) a curated brain metastasis marker panel derived from previously published signatures^7,33,95^; (2) label transfer from the Biermann et al. reference using *FindTransferAnchors* and *TransferData*; and (3) manual review of dot plots, violin plots, and spatial overlays.

Cells were assigned to three broad compartments (tumor, stroma, unknown). Each compartment was sub-clustered independently using the same Harmony workflow. Eleven stroma populations were identified and further sub-clustered at resolution 0.3 to resolve sub-states: T cells (9 sub-clusters), macrophages (12), myeloid/dendritic cells (12), neutrophils (8), fibroblasts (8), endothelial cells (9), pericytes (6), astrocytes/neurons (6), oligodendrocytes (2), plasma cells (4), and B cells (1). Brain parenchymal populations (neurons, astrocytes, oligodendrocytes) underwent a dedicated sub-clustering pass using k-nearest-neighbor label transfer in the full Harmony embedding. Twenty tumor sub-clusters were resolved in the tumor compartment; tumor cells were further classified by cancer of origin (melanoma, breast or unknown) from cancer-type-specific marker expression. A controlled-vocabulary status code (keep, atypical, mixed_lineage, unresolved_state, artifact_low_quality) was assigned to each cluster to enable flexible downstream filtering. Only artifact_low_quality clusters were excluded by default. Final labels were projected to all cells by k-nearest-neighbor transfer (RANN::nn2, k = 20) in the Harmony embedding.

#### Spatial Neighborhood Analysis

The spatial microenvironment of each cell was characterized by computing k = 12 nearest neighbors by Euclidean distance in tissue coordinates (µm) using RANN::nn2. The fractional composition of each neighborhood across 14 main cell-type categories was computed, yielding a cells × 14 neighborhood composition matrix per tissue core. Log2 enrichment scores were calculated as log2(observed / expected), where expected was the population-level frequency of each category within the same core. Spatial niches were identified by k-means clustering of the neighborhood composition matrix (k = 12, selected by elbow plot inspection, set.seed = 42).

#### GRN Myeloid Spatial Analysis

Granulin (GRN) expression was analyzed in the myeloid compartment of three samples (RD-23-1250, RD-25-171, RD-25-1585). Myeloid cells were stratified into two functionally distinct groups: Microglia (homeostatic, early activated, IFN-reactive, and DAM-like microglia) and Macrophage (anti-inflammatory BAM, meningeal BAMs, monocyte-derived TAMs, hypoxic TAMs and monocytes) by using FindMarkers() analysis (Table S7). GRN expression was normalized within each myeloid group; cells with detectable GRN expression (GRN_norm > 0) were stratified into GRN-high (≥ 80th percentile) and GRN-low (≤ 20th percentile) tiers, computed independently per myeloid group. GRN-mid cells were excluded from comparative analyses.

Two curated immune gene signatures were evaluated: broad immunosuppressive microenvironment, and antigen presentation (Table S8). Each signature was first restricted to genes present on the Xenium gene panel (5,089 genes total). For each cell, a signature score was calculated as the proportion of panel-detected signature genes with non-zero log-normalized expression (i.e., the fraction of the signature “on” in that cell). Scores were compared between *GRN*-high and *GRN*-low cells within each myeloid subclass using two-sided Wilcoxon rank-sum tests, with Benjamini-Hochberg correction applied across signatures. Notably, the entire classical MHC-I/II gene set is absent in the Human Xenium Panel (HLA-A, HLA-B, HLA-C, HLA-E, HLA-F, HLA-DRA, HLA-DRB1, HLA-DRB5, HLA-DQA1, HLA-DQA2, HLA-DQB1, HLA-DPA1, HLA-DPB1, HLA-DOA, HLA-DOB) together with CD74, TAPBP, B2M, and CALR; as a result, this signature on the Xenium panel captures co-stimulatory and accessory antigen-presentation machinery rather than MHC expression per se. Spatial neighborhood composition was assessed by two complementary approaches: (1) k = 100 nearest-cell neighborhoods (density-normalized, comparable across tissue regions) and (2) fixed-radius neighborhoods (50 µm; dbscan::frNN) representing physically defined proximity. Microglia and Macrophage groups were analyzed separately. Proximity to each T cell subtype was quantified as the distance to the nearest T cell of that subtype (RANN::nn2). The fraction of myeloid cells within 50 µm of exhausted CD8+ T cells (early, IFN, and terminally exhausted) was also compared between *GRN*-high and *GRN*-low groups using the Fisher exact test. A sign permutation test (N = 1,000; per-sample median distance; BH correction) tested whether *GRN*-high cells were systematically closer to each T cell subtype across samples. Pseudobulk summaries (per-sample mean) were computed for the 3-sample subset.

#### Software and Statistical Analysis

All analyses were performed in R v4.4.2 on the NYU Langone BigPurple/UltraViolet HPC. Key packages: Seurat v5, BPCells, harmony, RANN, dbscan, dplyr, ggplot2, patchwork, cowplot, rstatix, pheatmap. Differential expression and neighborhood composition differences were tested with the Wilcoxon rank-sum test; multiple testing was corrected using the Benjamini-Hochberg procedure. All stochastic steps used set.seed.

## Resource Availability

### Lead contact

Further information and requests for resources and reagents should be directed to and will be fulfilled by the lead contact, Eva Hernando.

### Material availability

Requests for resources and reagents should be directed to and will be fulfilled by the lead contact, Eva Hernando

### Data and code availability

All sequencing data will be deposited at [TBD] as [Database:TBD] and are publicly available as of the date of publication. Additional datasets utilized are detailed indicated in references, figure legends and Methods.

All code and packages are available in this paper’s methods.

Any additional information required to reanalyze the data reported in this paper is available from the lead contact upon request: Eva Hernando.

## Author Contributions

N.M.E., K.P.-C., J.W., and E.H. conceived and designed the study. N.M.E., S.A.A.S., and E.T.K. developed the brain-tropic tumor cell lines. N.M.E. collected tissue for mouse spatial transcriptomic profiling and performed the associated computational analyses. N.M.E., K.P.-C., and A.F.Y. performed all mouse experiments, cell culture, flow cytometry, and multiplex immunofluorescence experiments and analyses. L.H.G. assisted with BMDM culture experiments. M.H.G., M.U.G., and M.K. curated the human Visium cohort and data. M.H.G., M.U.G., M.K., G.B., A.S., and P.N. performed the human Visium analyses. D.A.S. performed neuropathologic review and histopathologic region classification of the human Visium specimens. M.A.G.-M., M.I., and I.I.B. performed the human Xenium analyses and human multiplex immunofluorescence experiments and analyses. N.M.E., K.P.-C., M.A.G.-M., and M.K. prepared figures and tables for this manuscript. H.J., A.L., M.H.G., J.W., and E.H. provided supervision and scientific guidance. N.M.E., K.P.-C., and E.H. wrote the manuscript. All authors reviewed and approved the manuscript.

## Acknowledgments

This work was supported by U54CA263001, R21CA286244, and R01CA277425 (E.H.), U54CA261717 (M. H.-G.), R37CA273333 (J.W.), R01CA269898 (J.W.) and R01AI190103 (J.W.) from NCI/NIH, the Melanoma Research Alliance (E.H.)(J.W.) and the V Foundation for Cancer Research (E.H.). N.M.E. was supported by F30CA288047 (NCI/NIH). K.P.-C. was supported by T32GM007308 (NIH).

We thank the Genomics Technology Center (GTC), particularly Paul Zappile MS, and Mr. Peter Meyn, for supervising all the sequencing; the Preclinical Imaging Core (Orlando Aristizabal MPhil), the Experimental Pathology Core (Director Dr. Cynthia Loomis, Gyles Ward MS, Shanmugapriya Selvaraj MS, Kalina Rice MS, Mr. Mark Alu, Adoni Dowridge MPH and Dr. Valeria Mezzano) and the Flow Cytometry core, all of which are supported by the Cancer Center Support Grant (CCSG) from NCI/NIH to the NYULH Perlmutter Cancer Center (P30CA016087; PI: Anirban Maitra). We thank Uta Mackensen for her beautiful work on our graphical abstract.

## Conflicts of interests

J.W. has sponsored research funds from Remunix in the past 12 months. J.W. has been a consultant or advisor for Remunix, Bristol Myers Squibb, Regeneron, Henlius, and Hanmi, and is the founder and equity holder for Remunix and Feedback Therapeutics. All other authors declare no conflicts of interest.

## Key Resources

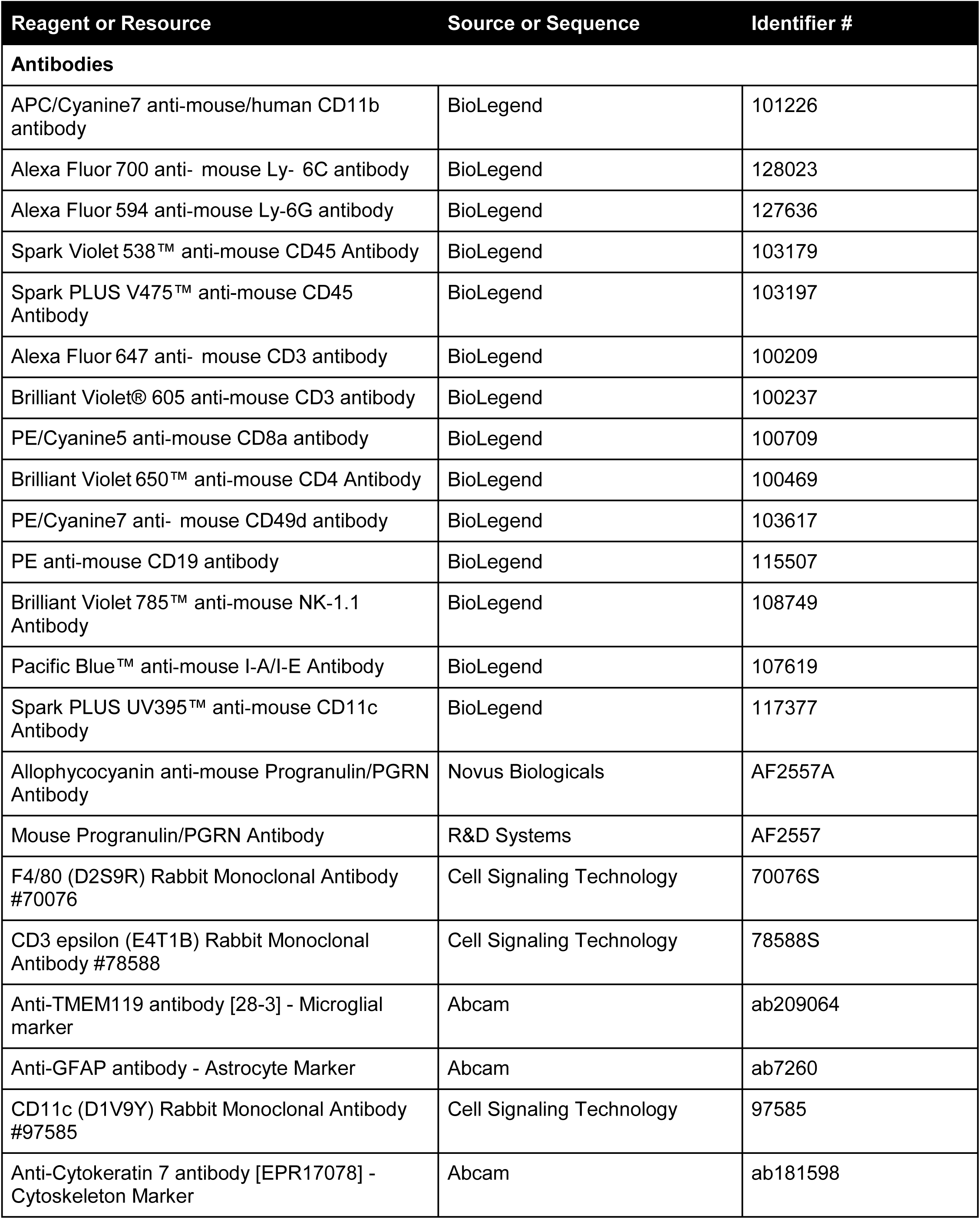
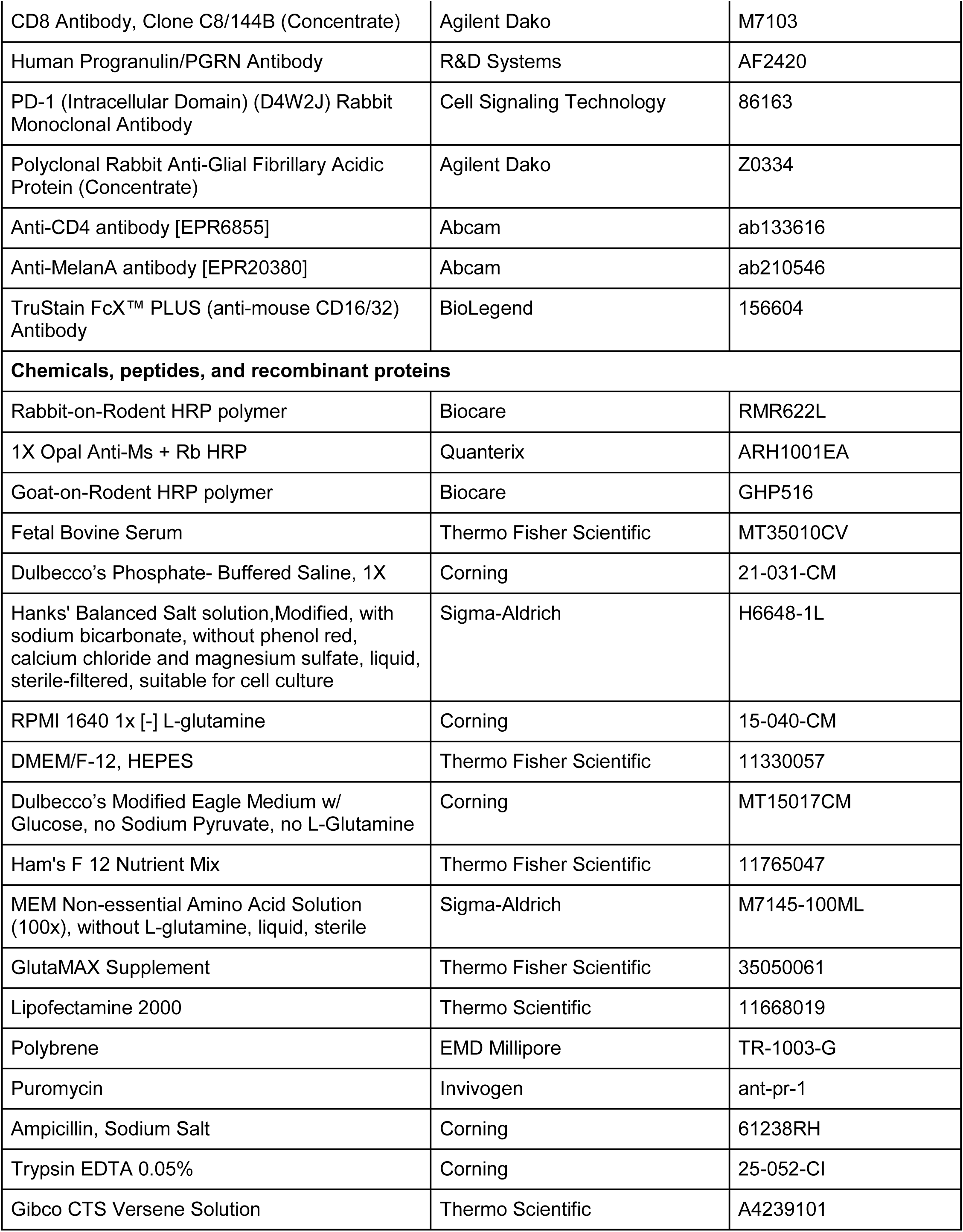
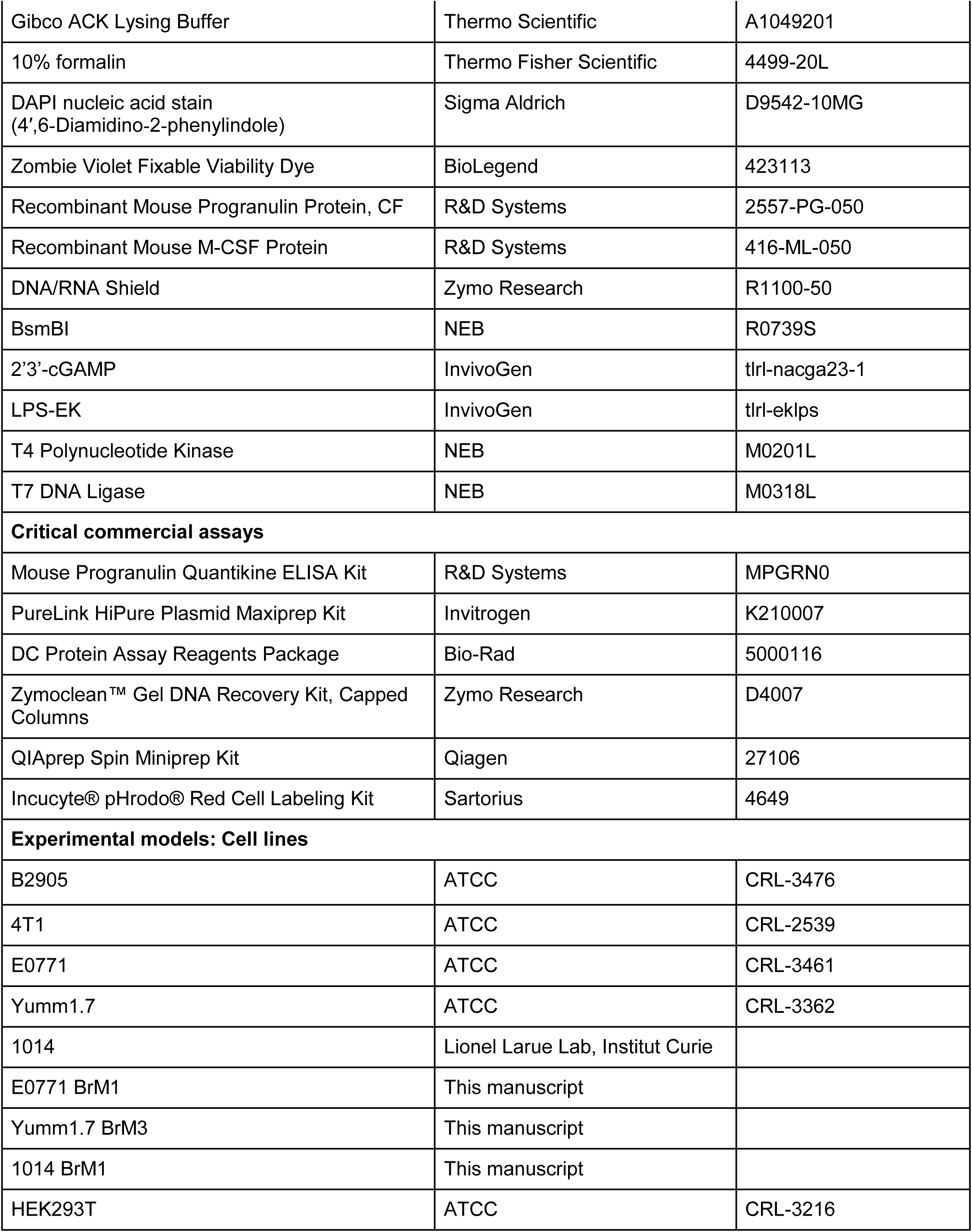
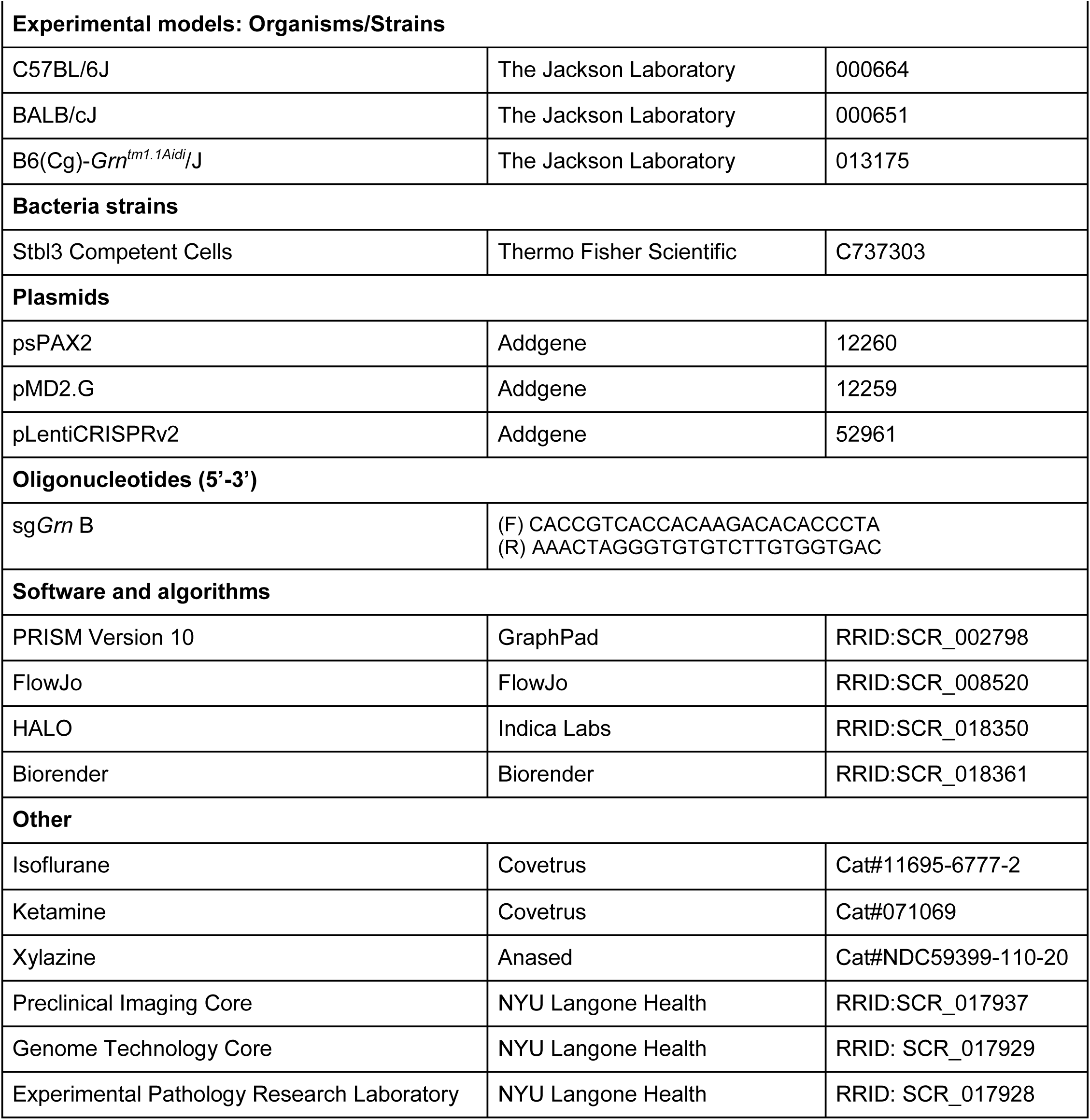

| R Packages |  |
| --- | --- |
| R package | Version |
| AnnotationDbi | 1.66.0 |
| Banksy | 1.0.0 |
| BayesSpace | 1.14.0 |
| Biobase | 2.64.0 |
| BiocGenerics | 0.50.0 |
| biomaRt | 2.60.1 |
| CellChat | 2.1.2 |
| circlize | 0.4.16 |
| cluster | 2.1.8.1 |
| clusterProfiler | 4.12.6 |
| ComplexHeatmap | 2.20.0 |
| cowplot | 1.1.3 |
| data.table | 1.17.0 |
| dplyr | 1.1.4 |
| enrichplot | 1.24.4 |
| forcats | 1.0.0 |
| GenomeInfoDb | 1.40.1 |
| GenomicRanges | 1.56.2 |
| ggcorrplot | 0.1.4.1 |
| ggplot2 | 3.5.2 |
| ggpubr | 0.6.0 |
| ggrepel | 0.9.6 |
| GOSemSim | 2.30.2 |
| harmony | 1.2.3 |
| igraph | 2.1.4 |
| IRanges | 2.38.1 |
| liana | 0.1.14 |
| lubridate | 1.9.4 |
| Matrix | 1.7-3 |
| MatrixGenerics | 1.16.0 |
| matrixStats | 1.5.0 |
| monocle3 | 1.3.7 |
| nicheDE | 0.0.0.9000 |
| nichenetr | 2.2.0 |
| NICHES | 1.1.0 |
| NMF | 0.28 |
| org.Hs.eg.db | 3.19.1 |
| org.Mm.eg.db | 3.19.1 |
| patchwork | 1.3.0 |
| pheatmap | 1.0.12 |
| presto | 1.0.0 |
| purrr | 1.0.4 |
| RColorBrewer | 1.1-3 |
| Rcpp | 1.0.14 |
| readr | 2.1.5 |
| registry | 0.5-1 |
| rlist | 0.4.6.2 |
| rngtools | 1.5.2 |
| rrvgo | 1.21.1 |
| S4Vectors | 0.42.1 |
| scales | 1.4.0 |
| scCustomize | 3.0.1 |
| sctransform | 0.4.2 |
| Seurat | 5.3.0 |
| SeuratObject | 5.1.0 |
| SeuratWrappers | 0.3.5 |
| SingleCellExperiment | 1.26.0 |
| sp | 2.2-0 |
| SPOTlight | 1.8.0 |
| STdeconvolve | 1.8.0 |
| stringr | 1.5.1 |
| SummarizedExperiment | 1.34.0 |
| tibble | 3.2.1 |
| tidyr | 1.3.1 |
| tidyverse | 2.0.0 |
| uwot | 0.2.3 |

